# Insights into the secondary structural ensembles of the full SARS-CoV-2 RNA genome in infected cells

**DOI:** 10.1101/2020.06.29.178343

**Authors:** Tammy C. T. Lan, Matthew F. Allan, Lauren E. Malsick, Stuti Khandwala, Sherry S. Y. Nyeo, Yu Sun, Junjie U. Guo, Mark Bathe, Anthony Griffiths, Silvi Rouskin

**Affiliations:** Whitehead Institute for Biomedical Research, Cambridge, Massachusetts, USA; Computational and Systems Biology, Massachusetts Institute of Technology, Cambridge, Massachusetts, USA; Department of Biological Engineering, Massachusetts Institute of Technology, Cambridge, Massachusetts, USA; National Emerging Infectious Diseases Laboratories, Boston University School of Medicine, Boston University, Boston, Massachusetts, USA; Department of Biology, Massachusetts Institute of Technology, Cambridge, Massachusetts, USA; Department of Electrical Engineering & Computer Science, Massachusetts Institute of Technology, Cambridge, Massachusetts, USA; Department of Neuroscience, Yale University School of Medicine, New Haven, CT, USA

## Abstract

SARS-CoV-2 is a betacoronavirus with a single-stranded, positive-sense, 30-kilobase RNA genome responsible for the ongoing COVID-19 pandemic. Currently, there are no antiviral drugs with proven efficacy, and development of these treatments are hampered by our limited understanding of the molecular and structural biology of the virus. Like many other RNA viruses, RNA structures in coronaviruses regulate gene expression and are crucial for viral replication. Although genome and transcriptome data were recently reported, there is to date little experimental data on native RNA structures in SARS-CoV-2 and most putative regulatory sequences are functionally uncharacterized. Here we report secondary structure ensembles of the entire SARS-CoV-2 genome in infected cells at single nucleotide resolution using dimethyl sulfate mutational profiling with sequencing (DMS-MaPseq) and the algorithm ‘detection of RNA folding ensembles using expectation–maximization’ clustering (DREEM). Our results reveal previously undescribed alternative RNA conformations across the genome, including structures of the frameshift stimulating element (FSE), a major drug target, that are drastically different from prevailing *in vitro* population average models. Importantly, we find that this structural ensemble promotes frameshifting rates (~40%) similar to *in vivo* ribosome profiling studies and much higher than the canonical minimal FSE (~20%). Overall, our result highlight the value of studying RNA folding in its native, dynamic and cellular context. The genomic structures detailed here lays the groundwork for coronavirus RNA biology and will guide the design of SARS-CoV-2 RNA-based therapeutics.

## Introduction

Severe acute respiratory syndrome coronavirus 2 (SARS-CoV-2) is the causative agent of coronavirus disease 2019 (COVID-19), recently declared a global pandemic by the World Health Organization (WHO). SARS-CoV-2 is an enveloped virus belonging to the genus *betacoronavirus*, which also includes SARS-CoV, the virus responsible for the 2003 SARS outbreak, and Middle East respiratory syndrome coronavirus (MERS-CoV), the virus responsible for the 2012 MERS outbreak. Despite the devastating effects these viruses have had on public health and the economy, currently no effective antiviral treatment exist. There is therefore an urgent need to understand their unique RNA biology and develop new therapeutics against this class of viruses.

Coronaviruses (CoVs) have single-stranded and positive-sense genomes that are the largest of all known RNA viruses (27 – 32 kb) (Masters, 2006). Prior to the emergence of SARS-CoV-2, most studies on secondary structures within coronavirus RNA genomes focused on several conserved regions that are essential viral replication: the 5’ UTR, the 3’ UTR, and the frameshift stimulating element (FSE) (Plant *et al.*, 2005; Yang and Leibowitz, 2015). Functional studies have revealed the importance of their secondary structures for viral transcription and replication (Brierley, Digard and Inglis, 1989; Liu *et al.*, 2007; Li *et al.*, 2008; Yang and Leibowitz, 2015).

The FSE straddles the boundary of ORF1a and ORF1b and causes the ribosome to “slip” and shift register by −1 nt in order to bypass a stop codon at the end of ORF1a and translate to the end of ORF1ab, producing a large polyprotein comprising 15 nonstructural proteins (nsps), including the viral RNA-dependent RNA polymerase (nsp12) and helicase (nsp13) (Brierley *et al.*, 1987; Plant *et al.*, 2005). Studies on multiple coronaviruses have shown that an optimal ribosomal frameshifting rate is critical, and small differences in percentage of frameshifting lead to dramatic differences in genomic RNA production and infection dose (Plant *et al.*, 2010). Therefore, the FSE has emerged as a major drug target for binding of small molecules that can influence the rate of ribosome slippage and is under active investigation to be used as a treatment against SARS-CoV-2 (Sun *et al.*, 2020; Zhang *et al.*, 2020).

The structures of coronavirus FSEs have been studied extensively. Short segments of the core FSE from both SARS-CoV-1 (Plant *et al.*, 2005) and SARS-CoV-2 (Zhang *et al.*, 2020) fold into complex structure with a three-stemmed pseudoknot. Small molecules, locked nucleic acids (LNAs), and mutations that are intended to disrupt this structure have been shown to impair viral replication (Kelly *et al.*, 2020; Sun *et al.*, 2020; Zhang *et al.*, 2020). However, despite the importance of the FSE structure, there is to date no direct validation of the relationship between the RNA folding conformation and frameshifting rate in infected cells.

Over the last decade, major advances in methods for RNA chemical probing have enabled genome-wide characterization of RNA structures in living cells. The most commonly used chemical probes are dimethyl sulfate (DMS) (Rouskin *et al.*, 2014) and reagents in the SHAPE (Siegfried *et al.*, 2014) and icSHAPE (Spitale *et al.*, 2015) families. DMS reacts with the Watson-Crick face of adenine (A) and cytosine (C) bases and probes base pairing directly, while SHAPE and icSHAPE reagents react with the 2’-OH group of all four nucleotides and measure nucleotide flexibility as a proxy for base pairing (Cordero *et al.*, 2012). Predictions of RNA structure that use DMS reactivities as folding constraints are of similar or marginally higher accuracies than predictions using SHAPE reactivities, as the specificity of DMS for Watson-Crick base-pairing compensates for the ability of SHAPE to probe all four nucleotides (Cordero *et al.*, 2012).

Two studies (Huston *et al.*, 2020; Manfredonia *et al.*, 2020) recently proposed models of the secondary structure of the entire genome of SARS-CoV-2 in Vero cells using SHAPE-MaP (Siegfried *et al.*, 2014). Both of these models are based on the average SHAPE reactivities at each nucleotide, so they cannot provide direct experimental evidence of alternative structures. However, the genomes of RNA viruses form not one structure but an ensemble of many structures whose dynamics regulate critical viral processes, such as splicing in HIV-1 (Tomezsko *et al.*, 2020). Thus, more work is needed to determine the dynamics of RNA structures within the SARS-CoV-2 genome and their functional roles in the viral life cycle.

In this study, we perform DMS mutational profiling with sequencing (DMS-MaPseq) (Zubradt *et al.*, 2016) and DREEM clustering (Tomezsko *et al.*, 2020) on SARS-CoV-2 infected Vero cells to generate the first insights into experimentally determined, single-nucleotide resolution genome-wide secondary structure ensembles of SARS-CoV-2. Our results reveal major differences with *in silico* and population-average structure predictions. Importantly, we highlight the physiological structure dynamics of known functional elements, such as the alternative structures at the FSE that determine frameshifting rates in cells. Our work provides experimental data on the structural biology of RNA viruses and will inform efforts on the development of RNA-based diagnostics and therapeutics for SARS-CoV-2.

## Results

### The genome-wide structure of SARS-CoV-2 in cells

To determine the intracellular genome-wide structure of SARS-CoV-2, we added dimethyl sulfate (DMS) to infected Vero cells and performed mutational profiling with sequencing (DMS-MapSeq) (Zubradt *et al.*, 2016) (Figure 1A). We chose DMS because it rapidly modifies unpaired adenines (A) and cytosines (C) *in vivo* at their Watson-Crick faces with negligible background effects (Zubradt *et al.*, 2016) and has been shown to yield structures of similar or slightly higher accuracies compared to SHAPE (Cordero *et al.*, 2012). Our results were highly reproducible between independent biological replicates (R^2^= 0.87; Figure 1B). Combined, a total of 87.2 million pairs of reads mapped to the coronavirus genome (Figure 1C), representing ~40% of total cellular RNA (post ribosomal RNA depletion). This large fraction of coronavirus reads from total intracellular RNA is consistent with previous literature using SARS-CoV-2 infected Vero cells (Kim *et al.*, 2020). DMS treated samples had high signal to noise ratio, with adenines and cytosines having a mutation rate ~9-fold higher than the background (guanines and uracils). In contrast, in untreated samples the mutation rate on all four bases (0.10%) was slightly lower than previously reported average sequencing error rates of 0.24% (Pfeiffer *et al.*, 2018) (Figure 1D). We used the DMS-MaPseq data as constraints in RNAstructure (Mathews, 2004) to fold the entire SARS-CoV-2 genomic RNA (Supplementary Figure 1) and assessed the quality of our model using two approaches.

**Figure 1:**
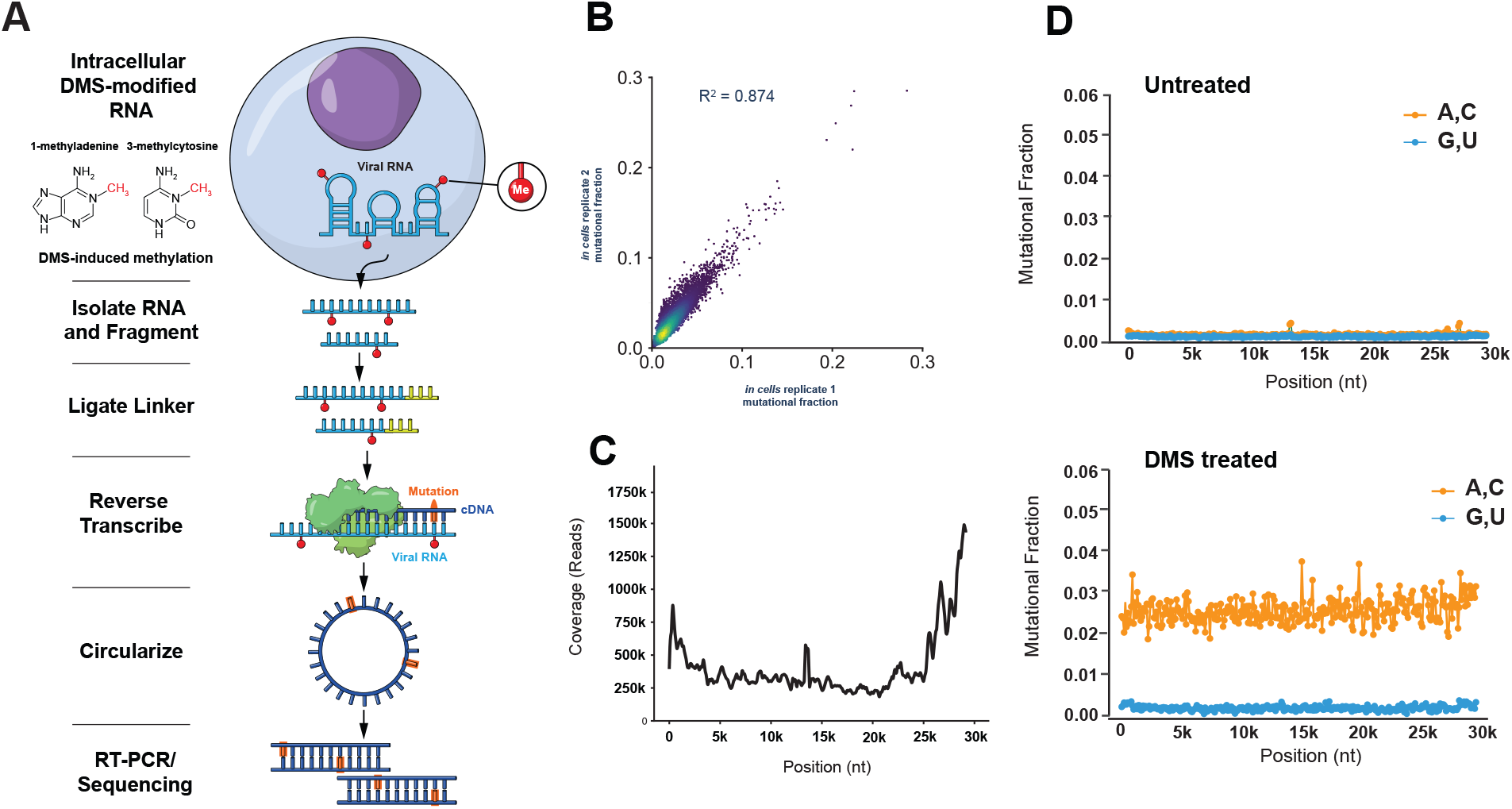
Genome-wide probing of intracellular SARS-CoV-2 RNA structure with DMS-MaPseq. **(A)** Schematic of the experimental protocol for probing viral RNA structures with DMS-MaPseq. **(B)** Correlation of DMS reactivities for each base between two biological replicates. **(C)** Genome-wide coverage as a function of position. Coverage at each position represents the average coverage over a 400 nt window. **(D)** Signal and noise as a function of genome position for untreated and DMS-treated RNA. Signal (mutation rate for A and C) and noise (mutation rate for G and U) at each position was plotted as the average of 100 nt window. Mutational Fraction of 0.01 at a given position represents 1% of reads having a mismatch or deletion at that position.

First, we introduce the data-structure correlation index (DSCI), a new metric based on the Mann-Whitney *U* statistic (Mann and Whitney, 1947) for quantifying how well a secondary structure model is supported by underlying chemical or enzymatic probing data (Figure 2A; see Methods). For probes that preferentially react with unpaired bases (e.g. DMS), the DSCI is defined as the probability that a randomly chosen unpaired base in the predicted structure will have higher reactivity than a randomly chosen paired base (for DMS, only A and C residues are considered for this calculation). A DSCI of 1 indicates perfect agreement between structure and data, 0.5 indicates no relationship, and 0 indicates complete disagreement.

**Figure 2:**
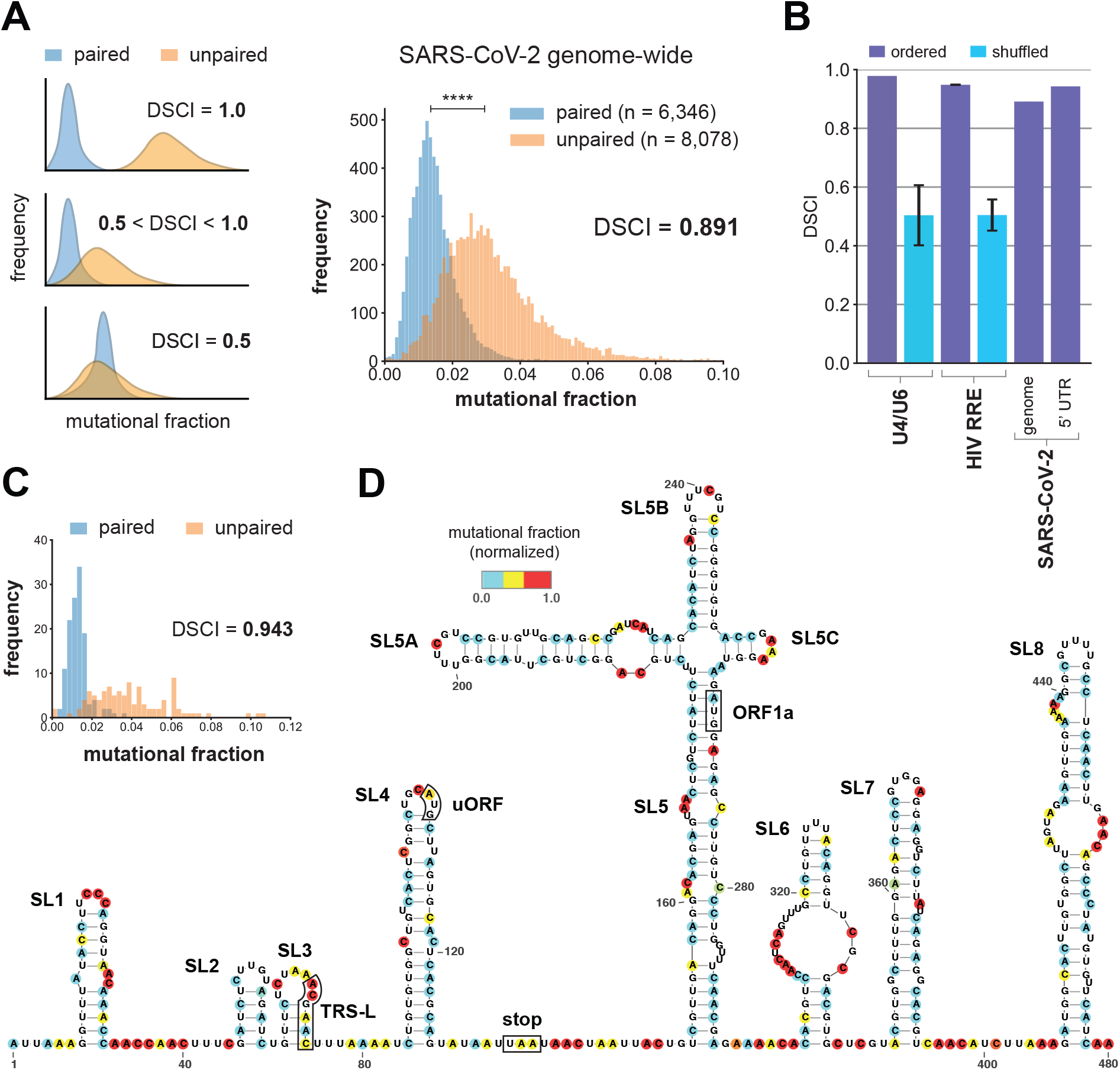
Quality assessment of the SARS-CoV-2 secondary structure model genome-wide and of the 5’ UTR. **(A)** (Left) Schematic of the data-structure correlation index (DSCI) showing possible distributions of signal on paired and unpaired bases for DSCI values of 1.0, 0.5, and intermediate. (Right) Distribution of signal on paired and unpaired bases genome-wide, and value of DSCI. The n = 55 bases (0.38% of all 14,424 As and Cs) with mutational fractions >0.10 are not shown for visual clarity, but are included in DSCI calculation. Horizontal bar indicates median values of paired and unpaired distributions, and **** indicates *P* < 0.0001 (Mann-Whitney *U* test). **(B)** DSCI values of control RNAs U4/U6 and HIV RRE, and of the whole SARS-CoV-2 genome and the 5’ UTR. Error bars (if present) show standard deviations between replicates. Shuffled negative controls show the distribution of DSCI when the signals on A and C residues were shuffled randomly 100 times. **(C)** Distribution of signal on paired and unpaired bases in the first 480 nt of the genome (structure in 1D), and value of DSCI. **(D)** In-cell model of the first 480 nt of the genome, including the 5’ UTR and sequences immediately downstream. Bases are colored by their DMS signal; bases that are not DMS reactive are colored white.

We confirmed that DSCI measures data-structure agreement using two RNAs with well-defined structures that we had previously analyzed with DMS-MaPseq (Tomezsko *et al.*, 2020): the U4/U6 snRNA and the HIV Rev Response Element (RRE) (Figure 2B). U4/U6 in vitro had a DSCI of 0.978 relative to its crystal structure (Cornilescu *et al.*, 2016), while RRE in cells had a DSCI of 0.949. As negative controls, we randomly shuffled the reactivities of all of A and C residues 100 times and computed the DSCI for each permutation; for each RNA, the mean DSCI for shuffled reactivities was approximately 0.50, as expected for random data. Thus, DSCI accurately measures how well the structure model is supported by the data, with DSCI values of roughly 0.95 or greater indicating very strong support.

Our genome-wide structure model was well-supported by our chemical probing data, with a global DSCI of 0.891 (Figure 2B) and significantly greater reactivities of unpaired bases relative to paired bases (*P* < 0.0001, Mann-Whitney *U* test; Figure 2A). We note that two previously published in-cell genome-wide models agreed substantially less with their respective chemical probing datasets, with global DSCI values of 0.705 (Huston *et al.*, 2020) and 0.760 (Manfredonia *et al.*, 2020) (Supplementary Figure 2).

Second, we found that our model of the 5’ untranslated region (UTR) agreed well with previous studies, showing that we could accurately identify known secondary structures (Figure 2D). The secondary structures of the 5’ UTR are conserved in multiple coronaviruses and have been characterized extensively (Yang and Leibowitz, 2015; Madhugiri *et al.*, 2018; Huston *et al.*, 2020; Manfredonia *et al.*, 2020; Miao *et al.*, 2020; Rangan, Zheludev and Das, 2020). In agreement with previous studies, we found five stem loops (SL1 – 5) within the 5’ UTR (nucleotides 1 – 265). These structures perform essential functions in viral replication (SL1 (Li *et al.*, 2008) and SL2 (Liu *et al.*, 2007)), subgenomic RNA production (SL3 (Yang and Leibowitz, 2015) and SL4 (Yang *et al.*, 2011)), and escape of nsp1-mediated translational suppression (SL1 (Banerjee *et al.*, 2020)). SL5 contains the start codon of ORF1 and branches into three additional stems (SL5A, SL5B, SL5C), a complex structure that our model recapitulates perfectly with respect to previous studies (Huston *et al.*, 2020; Miao *et al.*, 2020).

A short stem loop (SL4.5) has been proposed to occur between SL4 and SL5 (Huston *et al.*, 2020; Miao *et al.*, 2020; Rangan, Zheludev and Das, 2020). Our data suggest that SL4.5 does not exist, in agreement with another model based on in-cell data (Manfredonia *et al.*, 2020). Additional structures exist immediately downstream of the 5’ UTR. We found three stem loops (SL6 – 8) in this region, in nearly perfect agreement with two previous in-cell studies (Huston *et al.*, 2020; Manfredonia *et al.*, 2020). Further supporting the accuracy of our model, the DSCI was 0.943 across SL1 – 8, indicating that our model of this region agrees very well with our chemical probing data (Figure 2C).

### Genome structures that are well supported by multiple lines of evidence

To identify structures within the genome that are well supported by multiple types of evidence, we compared several types of evidence.

We compared our structure to two other genome-wide models of the SARS-CoV-2 genome structure in Vero E6 cells: Model 1 (Huston *et al.*, 2020) and Model 2 (Manfredonia *et al.*, 2020). Relative to each other, the viral genomes in these three studies contain zero indels and mismatches at only seven positions. As a similarity metric, we introduce a modified version of the the Fowlkes-Mallowes index (mFMI) that measures agreement of base pairs and unpaired bases (see Methods). Globally, our model was 81.4% similar to Model 1 and 80.7% similar to Model 2, while Models 1 and 2 were 76.4% similar.

To determine local similarity, we computed the mFMI across the genome using a sliding window of 80nt and a step size of 1nt (Supplementary Figure 3).

We evaluated the robustness of our in-cell data derived genome-wide model by varying two critical RNA folding parameters used by RNAstructure: 1) the maximum allowed distance for base pairing and 2) the threshold for DMS signal normalization.

A previous *in silico* approach for folding RNA found that limiting base pairs to be 100 to 150 nt apart was optimal to avoid overpredicting structured regions (Lange *et al.*, 2012). However, some RNA viruses contain known essential structures wherein bases over 300 nt apart are paired (e.g. the Rev response element in HIV-1 spans approximately 350 nt (Watts *et al.*, 2009)). We therefore varied the maximum distance (md) allowed for base pairing from 120 nt to 350 nt. We computed the agreement between the resulting structures using a modified version of the Fowlkes-Mallows index (Fowlkes and Mallows, 1983) that compares base pairing partners as well as unpaired bases (Methods). Overall, there was high agreement while varying the md from 120 nt to 350 nt, suggesting that long-distance (i.e. >120 nt) interactions across the SARS-CoV-2 RNA have a small effect on the identity of local structures. The genome structure folded with an md of 120 nt was 97.5% identical to the structure with an md of 350 nt, and in the latter structure only 3.8% of base pairs spanned >120 nt. Next, we proceeded with the md limit of 350 nt and tested two different DMS signal thresholds that normalize reactivity to either the median of the top 5 % or top 10% of the most reactive bases. We found that the structure models produced with the two normalization approaches were highly similar, with 93.6% identity (Figure 3A). Thus, within the ranges that we tested, our genome-wide data-derived model was robust to variation in the parameters of RNAstructure (Mathews, 2004).

**Figure 3:**
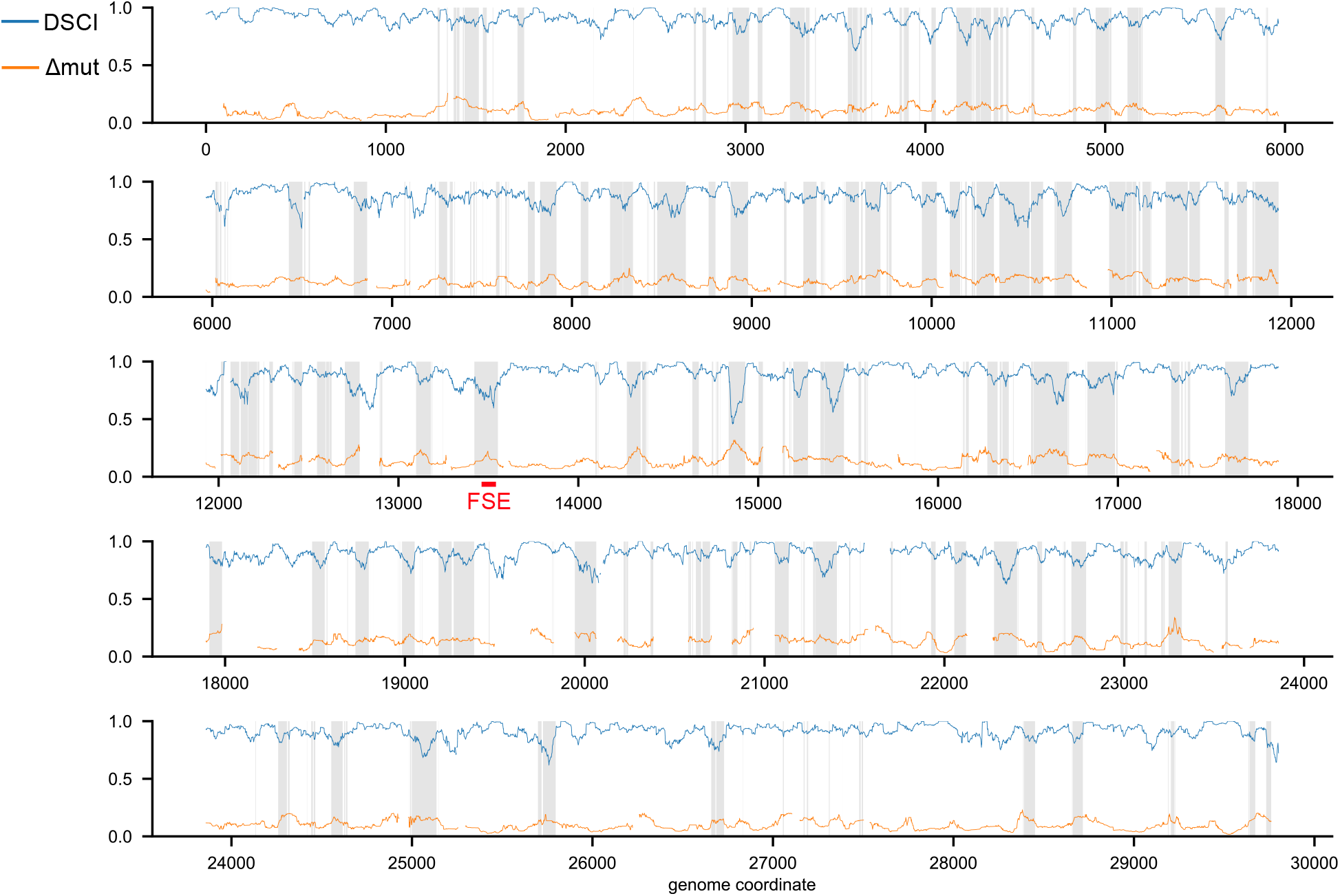
Genome-wide data-structure correlation index (DSCI) and inter-cluster change in DMS reactivity (ΔDMS) DSCI is computed for all overlapping 80nt windows genome-wide, except windows with fewer than 5 unpaired and 5 paired bases. ΔDMS is computed as the moving median for all overlapping 80nt windows containing at least 10 bases with DMS reactivities, after removing cluster regions with fewer than 100,000 reads or one cluster with a DMS reactivity greater than 0.3. Regions where clustering is likely to improve structure predictions over the population average model (with DSCI < 0.902 and ΔDMS > 0.117) are shaded in gray. The location of the frameshift stimulating element (FSE) is highlighted in red.

We proceeded with the whole genome structure modelled with a md of 350 nt and a DMS signal normalization of 5% for further analysis. Previous studies that computationally predicted genome-wide SARS-Cov-2 RNA structures used 1) RNAz, a thermodynamic-based model that additionally takes sequence alignment and considers base pairing conservation (Gruber *et al.*, 2010; Rangan, Zheludev and Das, 2020), and 2) Contrafold, which predicts RNA secondary structures without physics-based models and instead uses learned parameters based on known structures (Do, Woods and Batzoglou, 2006). These recent studies predicted 228 structures with RNAz with lengths ranging from 90 to 120 nt, and 79 structures with Contrafold with lengths ranging from 55 to 111 nt (Rangan, Zheludev and Das, 2020). For each of these structures, we computed the agreement between the different models (Supplementary Figure 4B). We report the agreement using the mFMI while either excluding external bases pairs or including these pairs (Methods). As expected, agreement with the structures from purely computational prediction is higher when excluding external base pairs (average 76.3% for RNAz, 69.3% for Contrafold) than when including them (average 71.2% for RNAz, 54.0% for Contrafold). Since our goal is to compare the overall similarity of two structures, we chose the inclusion of external base pairs as the more accurate metric for comparing the structures. Our predictions overall agreed more with those from RNAz (mean 71.2%, median 75.2%) than Contrafold (mean 54.0%, median 54.4%). We report the agreement between our structure and the RNAz structures across the entire genome (Figure 3C). Most structures are 60 to 80% identical, with several short regions that disagree substantially.

In addition, we computed the similarity of our model compared to the structures with the three highest P-values predicted with RNAz that do not overlap known structures in the Rfam database (Kalvari *et al.*, 2018; Rangan, Zheludev and Das, 2020) (Supplementary Figure 4D). We noted that in all three cases, the structure at the center of the window was nearly identical to ours, and most of the disagreements arose at the edges, presumably due to the effects of the windows from RNAz being limited to 120 nt. Of the five structures predicted with Contrafold that had the largest maximum expected accuracies, our agreement ranged from 66.0% to 86.1%, well above the genome-wide mean (54.0%), suggesting that these structures are indeed more accurate than the average Contrafold structure (Supplementary Figure 4E).

Finally, we compared the structures at the TRS elements to those predicted by RNAz (Rangan, Zheludev and Das, 2020) (Supplementary Figure 4F). To remove the effects of external base pairs, we focused only on the complete structural element (e.g. a stem loop) in which the TRS was located. RNAz predicted structure for four TRSs. Our model for TRS-L was identical to the first predicted window from RNAz but differed significantly (35.3% agreement) from the next prediction of the same TRS-L element within a different folding window, indicating that the choice of folding window can have a large effect on the RNAz structure model. For the other three TRS elements for which RNAz predicted at least one structure for, our agreement ranges from 74.4% to 96.8%, above the genome-wide average of 71.2%, lending support to both models.

To facilitate future studies investigating the binding of locked nucleic acid (LNA) probes to the genome, we determined the locations of all stretches of at least 14 consecutive unpaired bases in our genome-wide model (Supplementary Table 1). These 259 regions had a median length of 19 nt and a maximum length of 180 nt (at positions 21573 – 21752). Due to formation of alternative structures (discussed below), some of these regions may sometimes form base pairs, but they appear to exist at least some of the time in an unfolded state.

### Transcription-Regulating Sequences (TRSs) lie within stem loops

As the transcription-regulating sequences (TRSs) are necessary for the synthesis of sgRNAs, we analyzed our structural models of the leader TRS (TRS-L) and the nine body TRSs (TRS-B). The leader TRS (TRS-L) is the central component of the 5’ UTR involved in discontinuous transcription (Sola *et al.*, 2015). *In silico* models for several alpha and betacoronaviruses variously place TRS-L in stem loop 3 (SL3) or in an unpaired stretch of nucleotides (Liu *et al.*, 2007; Yang and Leibowitz, 2015). The TRS-L of SARS-CoV and of SARS-CoV-2 was predicted to lie in the 3’ side of the stem of SL3, which is consistent with our in-cell model (Liu *et al.*, 2007; Yang and Leibowitz, 2015; Rangan, Zheludev and Das, 2020). In our data, the stem of SL3 contains two bases with medium reactivity (Figure 2B), which suggests that SL3 transitions between folded and unfolded states, as is hypothesized for the alphacoronavirus transmissible gastroenteritis virus (TGEV) (Madhugiri *et al.*, 2018).

Of the nine body TRSs, we find that seven (all but the TRSs of ORF7a and ORF7b) lie within a stem loop. Of these, all but one TRS (N) place the core sequence on the 5’ side of the stem. Four body TRSs (M, ORF6, ORF8, and N) are predicted to lie in stem loops with two or three bulges, with the core sequence spanning one of the internal bulges. The other three structured body TRSs (S, ORF3a, and E) lie in stem loops without bulges, with the final paired base in the 5’ side of the stem contained in the core sequence. Strikingly, the entire core sequence is paired in two body TRSs (S and M), and partially exposed in a loop or bulge in the other five (Supplementary Figure 5B).

### The vast majority of the SARS-CoV-2 genome forms alternative structures

We previously discovered that for another ssRNA virus, HIV-1, over 90% of the genome forms ensembles of alternative structures rather than a single structure (Tomezsko *et al.*, 2020). Formation of alternative RNA structures has important functional consequences: for example, in HIV-1, they regulate alternative splicing. However, all previous studies that chemically probed the entire SARS-CoV-2 genome (Huston *et al.*, 2020; Manfredonia *et al.*, 2020) used only the average reactivity of each base to fold their structural models, and thus could not detect subpopulations of RNAs with different structures. Although these studies used Shannon entropy to estimate structural heterogeneity in a series of short sliding windows, this metric is still based on the average SHAPE reactivities per base and does not identify subpopulations of alternative structures directly from single molecule data.

We detected alternative structures in SARS-CoV-2 by applying the DREEM algorithm (Tomezsko *et al.*, 2020) to our in-cell DMS-MaPseq data. Briefly, DREEM clusters the sequencing reads based on which bases are DMS modified together on the same read and identifies sub-populations of molecules with distinct patterns of DMS modifications.

We partitioned the genome into 373 regions of 80 nt and clustered the reads mapping to each segment. All 316 regions that passed our quality control criteria (see Methods) formed at least two clusters, providing the first experimental evidence that the vast majority (at least 84%) of the SARS-CoV-2 genome forms alternative structures.

We hypothesized that if a region forms two very different alternative structures, the local agreement between the DMS reactivity data and the population average model (i.e. the DSCI) would be low, and vice versa. We computed the DSCI across the entire genome in overlapping windows of 80 nt with a step size of 1 nt (excluding windows without at least 5 paired and 5 unpaired bases with DMS reactivities). To quantify differences between alternative structures, we computed for each base in the region the difference in DMS mutation rate (hereafter ΔDMS) between the two clusters in each region (see Methods). In order to compare ΔDMS (a property of a single base) with DSCI (a property of multiple bases), we computed the moving median ΔDMS using the same sliding window as for DSCI (excluding windows with fewer than 10 bases with DMS reactivities).

Consistent with our hypothesis, DSCI and ΔDMS correlated negatively (r = −0.330, n = 26704, *P* < 0.0001), albeit weakly, indicating that large differences between alternative structures are associated with lower agreement between the population average structure and the DMS reactivities (Supplementary Figure 6). In support of the quality of our genome-wide model based on population average, there were no low-quality regions with minimal alternative structure (low DSCI and low ΔDMS). Surprisingly, many regions in the genome-wide model agreed well with the population average data, yet separated into clusters with large ΔDMS (high DSCI, high ΔDMS). For example, the region 14,561 – 14,640 folds into a bulged stem loop that is extremely well supported by the DMS reactivities (DSCI = 0.997) but forms distinct clusters of reactivity patterns, as indicated by the moderate ΔDMS of 0.105. We find that the reactivities of nucleotides in loops change considerably between the clusters, while the reactivities of nucleotides in stems change minimally. Thus, the set of nucleotides that are paired is mostly identical between the two clusters, while other factors cause changes in the reactivities of the nucleotides in loops, such as possibly changes in tertiary structure or transient formation of long-range RNA-RNA interactions. This finding indicates that predicted structures that agree strongly with the population average data are likely to be accurate, even if the data separate into distinct clusters due to changes in reactivities within loops.

In order to identify regions of the genome that did not correspond well to the population average model (low DSCI) and could be improved by clustering (high ΔDMS), we located all regions where the DSCI and ΔDMS were, respectively, below and above their genome-wide medians of 0.902 and 0.117 (Figure 3). For example, the genome-wide minimum DSCI (0.457) falls within the clustered region 14,881 – 14,960 and coincides with a peak in ΔDMS. We find that this region clusters into two distinct structural states: the major cluster (~80%) has an even distribution of DMS signal, suggesting unfolded or highly dynamic state; while the minor cluster (~20%) has an uneven distribution of DMS signal, suggesting a structured state. The structured state contains a stem loop spanning the same nucleotides (14,883 – 14,930) as a stem loop in the population average model, but the distal portion is considerably different and is much more consistent with the DMS reactivities of the structured state than the population average model is with the average reactivities. Thus, we find that clustering can identify secondary structures better supported by chemical probing data in regions where the population average model fails to generate a well-supported structure.

Interestingly, we find that the frameshift stimulating element (FSE), which is critical for regulating the translation of ORF1b, also has a structure that is poorly supported by the population average DMS reactivities and a large difference between clusters (Figure 3), suggesting that this region also forms multiple distinct structures. Although other studies have suggested that the FSE forms multiple structures, they have either inferred them indirectly using suboptimal folding based on population average reactivities (Huston *et al.*, 2020) or measured them in short segments of the FSE in vitro, outside of the context of genomic RNA and cellular factors (Neupane *et al.*, 2020). We find that the FSE indeed forms at least two distinct structures and characterize them in detail below.

### Uncovering an unexpected structure at Frameshift stimulating element (FSE)

The frameshift stimulating element (FSE) causes the ribosome to slip and shift register by −1 nt in order to bypass a stop codon and translate ORF1b, which encodes five non-structural proteins (nsps) including nsp12, an RNA-dependent RNA polymerase (RdRP) (Plant and Dinman, 2008). Previous studies on coronaviruses and other viruses have shown that an optimal frameshifting rate is critical and small differences in percentage of frameshifting lead to dramatic differences in genomic RNA production and infection dose (Plant *et al.*, 2010). Therefore, the FSE has emerged as a major drug target for small molecule binding that could influence the rate of frameshifting and be used as a treatment against SARS-CoV-2. To date, there is little experimental data on the structure of SARS-CoV-2 FSE and the prevailing model is a 3-stem pseudoknot forming downstream of the slippery site, which is thought to pause the ribosome and allow frameshifting to occur (Plant and Dinman, 2008).

To closely examine the FSE structure in cells, we used DMS-MaPseq target specific protocol (Zubradt *et al.*, 2016). We designed primers targeting 283 nt surrounding the FSE and amplified this region from cells infected with SARS-CoV-2 that were treated with DMS. Our analysis revealed a strikingly different structure than the prevailing model (Plant *et al.*, 2005; Rangan, Zheludev and Das, 2020) (Figure 4A). Our in-cell model does not include the expected pseudoknot formation downstream of the slippery sequence. Instead, half of the canonical stem 1 (Figure 4A, purple) finds an alternative pairing partner (pink) driven by 10 complementary bases upstream of the slippery site (Figure 4A, pink). We call this pairing Alternative Stem 1 (AS1).

**Figure 4:**
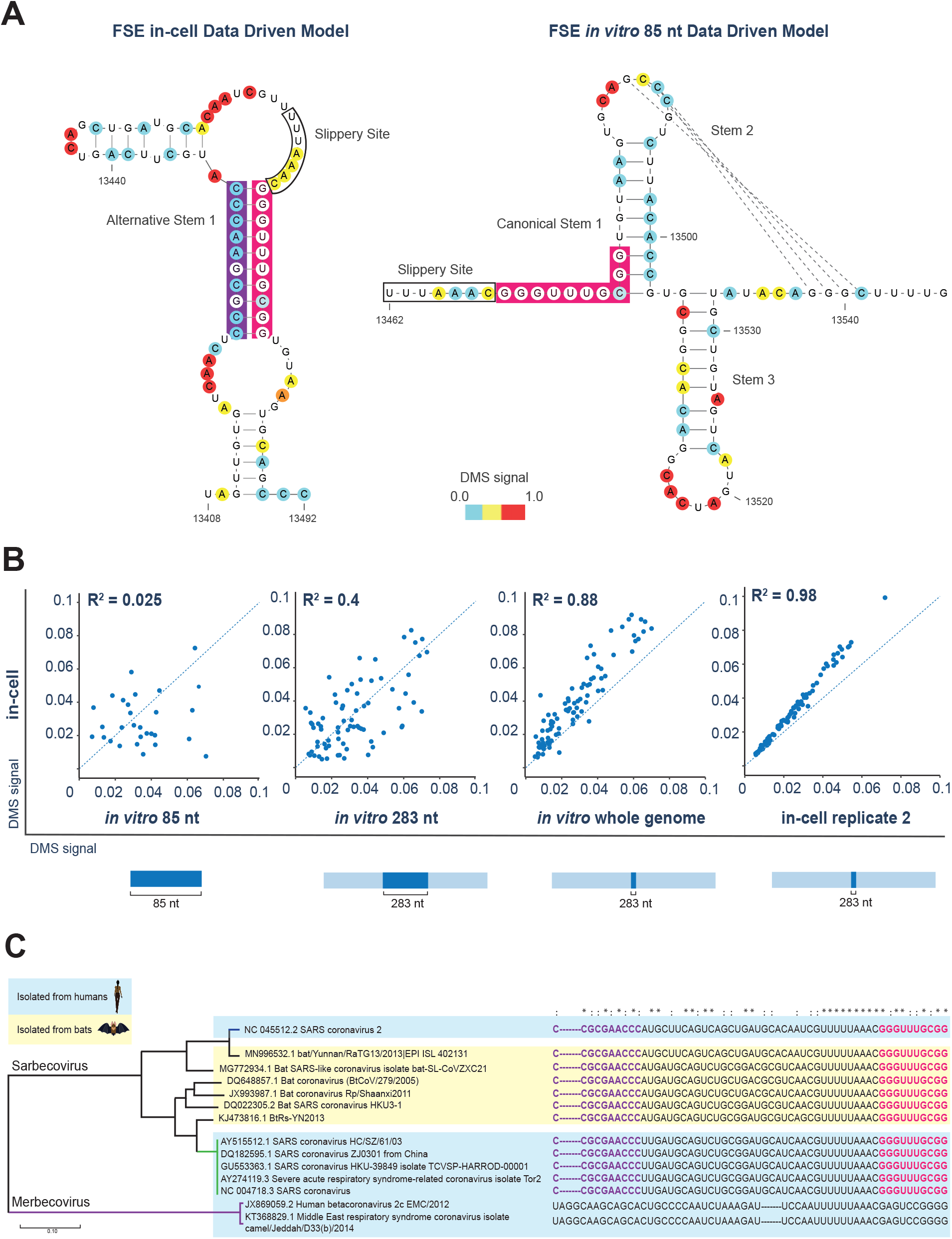
Frameshift stimulating element (FSE) adopts an unexpected structure in cells. **(A)** Structural model of FSE derived from DMS-MaPseq from (left) infected VERO cells and (right) *in vitro*-transcribed RNA. Nucleotides are color-coded by normalized DMS signal. The 5’ of the canonical stem 1 is highlighted in pink, the complement pairing (starting 46nt downstream) is shown in purple and the slippery site boxed in white. Structural model of *in vitro*-transcribed 85 nt FSE shown is the major cluster structure from DREEM clustering. **(B)** Scatter plots comparing FSE structures in different contexts. Comparison of DMS signals of in-cell replicate 1 with (leftmost) *in vitro* refolded 85 nt., (middle-left) *in vitro* refolded 283 nt., (middle-right) *in vitro* refolded whole genome, and (rightmost) in-cell replicate 2. The blue dotted line is the identity line; R is Pearson’s coefficient. **(C)** Sequence conservation of FSE alternative pairing. The 5’ sequence of canonical stem 1 is highlighted in pink and the complement sequence is highlighted in purple. Symbols above the sequences indicate perfect conservation among all viruses in the alignment (*) or perfect conservation among only the sarbecoviruses (:).

The prevailing model of the SARS-CoV-2 FSE is based on previous studies of the SARS-CoV FSE, as they only differ in sequence by a single nucleotide located in a putative loop (Rangan, Zheludev and Das, 2020). Nuclease mapping and Nuclear Magnetic Resonance (NMR) analysis of the SARS-CoV FSE solved the structure of an *in vitro* refolded, truncated 85 nt region starting at the slippery site (Plant *et al.*, 2005). This structure did not include the sequence upstream of the slippery site and formed a 3-stem pseudoknot.

Interestingly, *in silico* predictions of the RNA structure of the SARS-CoV-2 genome using RNAz (Rangan, Zheludev and Das, 2020) and ScanFold (Andrews *et al.*, 2020) do not find the 3-stem pseudoknot but instead support our in-cell model of Alternative Stem 1. In SARS-CoV-2, ScanFold not only predicted the AS1 but also found that it was more stable relative to random sequences than any other structure in the entire frameshift stimulating element (Andrews *et al.*, 2020). Indeed, three conceptually varied methods (DMS-MaPseq, RNAz, and ScanFold) aimed at identifying functional structures, run independently by different research groups all converge on the Alternative Stem 1 as a central structure at the FSE.

In order to directly compare our in-cell findings with the reports of the 3-stem pseudoknot, we *in vitro*-transcribed, refolded, and DMS-probed the same 92 nt sequence as analyzed by NMR (Plant *et al.*, 2005). Our in vitro-data driven model for the major cluster agrees well with the NMR model (87.1% identical) and finds all three canonical stems, including the pseudoknot.

### FSE structure is dependent on the sequence context

The major differences we observed in the structure of the FSE in cells vs. *in vitro* could either be due to 1) length of the *in vitro* refolded viral RNA or 2) factors in the cellular environment that are absent *in vitro*. To distinguish between these two possibilities, we re-folded the FSE in the context of longer native sequences.

We found that as we increased the length of the *in vitro* re-folded construct by including more of its native sequence, from 92 nt to 283 nt to 30 kb, the DMS reactivity patterns became progressively more similar to the pattern we observed in cells (Figure 4B). Indeed, in the context of the full ~30 kb genomic RNA, the structure of the FSE is nearly identical to the structure in physiological conditions during SARS-CoV-2 infection in cells (R^2^= 0.88). These results indicate that the length of the entire RNA molecule is important for correctly folding the FSE. Strikingly, at a length of 283 nt and above, the main structure forming is Alternative Stem 1 rather than the 3-stem pseudoknot. Our data indicate that given the full range of pairing possibilities in the genome, AS1 is more favorable and the predominant structure in cells.

### Alternative Stem 1 pairing sequence is conserved across sarbecoviruses

To determine if other coronaviruses may have a similar alternative structure of the frameshift stimulating element, we searched for the sequence that pairs with canonical stem 1 in a set of curated coronaviruses (Ceraolo and Giorgi, 2020). This set contains 53 isolates of SARS-CoV-2, 12 other sarbecoviruses (including the SARS-CoV reference genome), and 2 merbecoviruses. The 10 nt complement (CCGCGAACCC) to a sequence overlapping canonical stem 1 of the FSE (GGGUUUGCGG) was perfectly conserved in all 12 of the sarbecoviruses, six of which were isolated from bats (Figure 4C). However, the 10 nt complement was not present in either merbecovirus. Aligning the sequences of all 20 betacoronaviruses with complete genomes in RefSeq revealed that the 10 nt complement was conserved in all of and only the three sarbecoviruses in RefSeq: SARS-CoV, SARS-CoV-2, and BtCoV BM48-31 (data not shown). These results suggest that AS1 is unique to the sarbecoviruses.

### The Frameshift stimulating element (FSE) forms alternative structures in cells

We further analyzed the intracellular folding of the FSE using DREEM. We found two distinct patterns of DMS reactivities (Figure 5A), showing that the RNA folds into at least two distinct conformations at this region. In both biological replicates, Clusters 1 and 2 separate at a reproducible ratio (~54% vs. 46%) where Cluster 1 is drastically different from Cluster 2 (R^2^ = 0.25) but identical to the corresponding cluster in biological replicates (R^2^ = 0.99) (Figure 5B). Both structures have the Alternative Stem 1 pairing spanning the slippery sequence. As the pseudo-energy constrains for the DMS signal did not generate alternative structure models with high DSCI scores, we color coded the dynamic bases that change DMS signal between the two conformations onto the population average model (green and yellow, Figure 5B). Together with our *in vitro* results, which revealed that the folding at the FSE is influenced by the longer sequence context (Figure 4B), this data implies that the alternative conformations are driven by long-distance RNA:RNA interactions.

**Figure 5:**
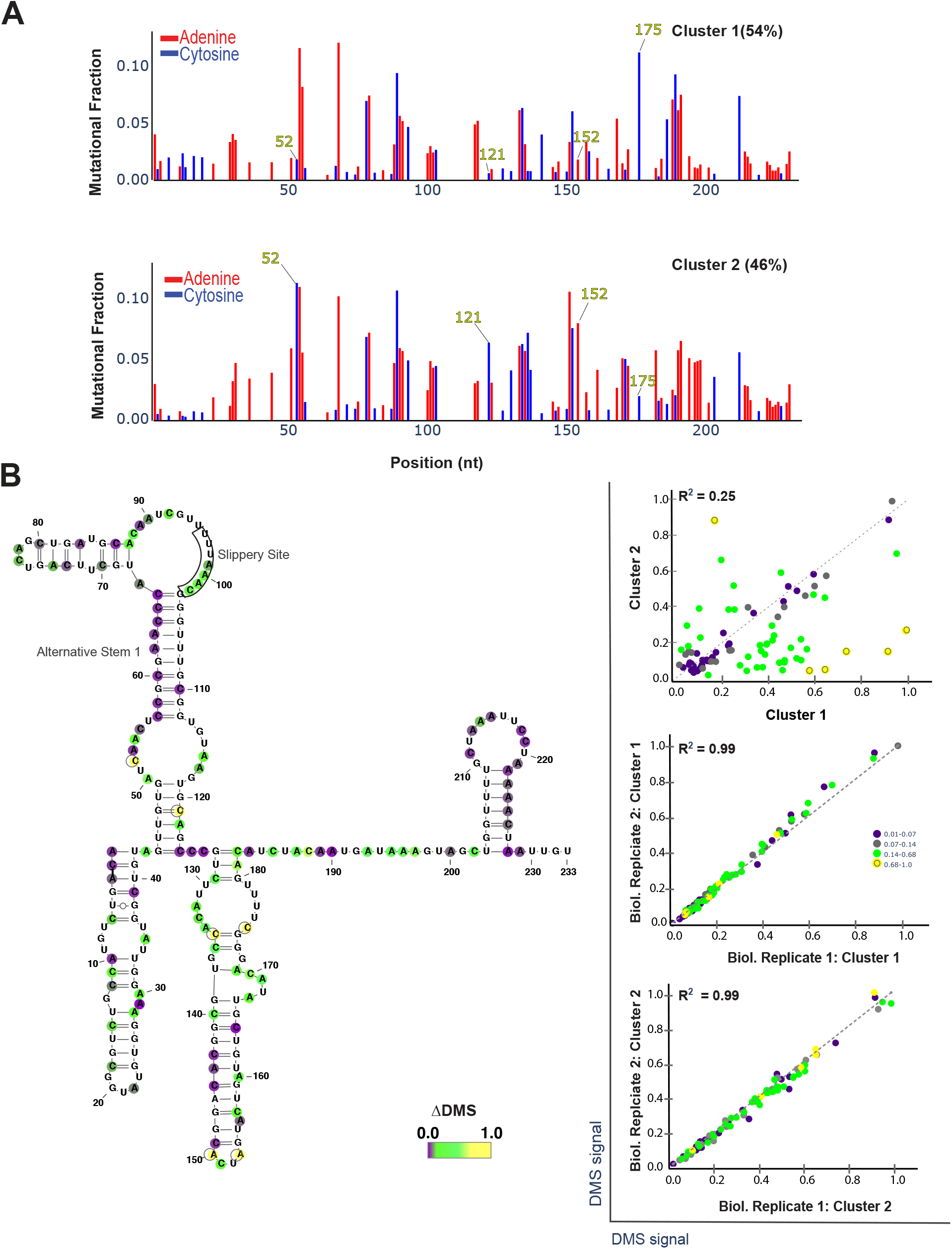
Alternative conformations of the frameshift stimulating element (FSE) derived from in-cell DMS-MaPseq data. **(A)** Clustered DMS signal for 283 nt surrounding the FSE structure from infected Vero cells, identified by DREEM clustering. Percentages for each cluster are determined by DREEM from representative sample of n = 2. **(B)** RNA structure dynamics of the FSE. (Left) Population average structure of the FSE. Alternative stem 1 and slippery site sequences are marked. Nucleotides are colored by change in DMS reactivity (ΔDMS). The same colors are used on the (Right) scatterplots showing the comparison of DMS signal between clusters; (Top) scatter plots of DMS signal between cluster 1 and 2 within a biological replicate; (Middle, Bottom) scatter plot of the variation in DMS signal for the same cluster between two biological replicates. The dotted line is the identity line; R is Pearson’s coefficient. The ΔDMS is the normalized distance of each point (i.e. nucleotide) to the identity line.

### Frameshifting rate is determined by FSE sequence context and structure

To directly measure how the FSE structure ensemble impacts frameshifting rate in cells, we constructed dual luciferase frameshift reporter constructs (Grentzmann *et al.*, 1998) with either a “short” FSE of only the 92nt region that folds into the canonical three-stemmed pseudoknot or a “long” FSE of the pseudoknot placed in the middle of approximately 3000nt of its native sequence context (Fig. 4a). The dual luciferase reporter is a well-established tool for measuring frameshifting rate, where the stop codon of a firefly luciferase (F-Luc) coding sequence is replaced with a FSE which allows a renilla luciferase (R-Luc) coding sequence in the −1 frame behind the FSE to report on frameshifting rate (Figure 6A). In addition, we *in vitro*-transcribed and transfected the reporter mRNA into cells to avoid cryptic transcription start sites or unintended splicing events of the DNA reporter that could impact F-Luc and R-Luc luminescence (Figure 6B). We calculated the frameshifting rate as the relative R-Luc to F-Luc ratio after normalization against negative and positive controls.

**Figure 6:**
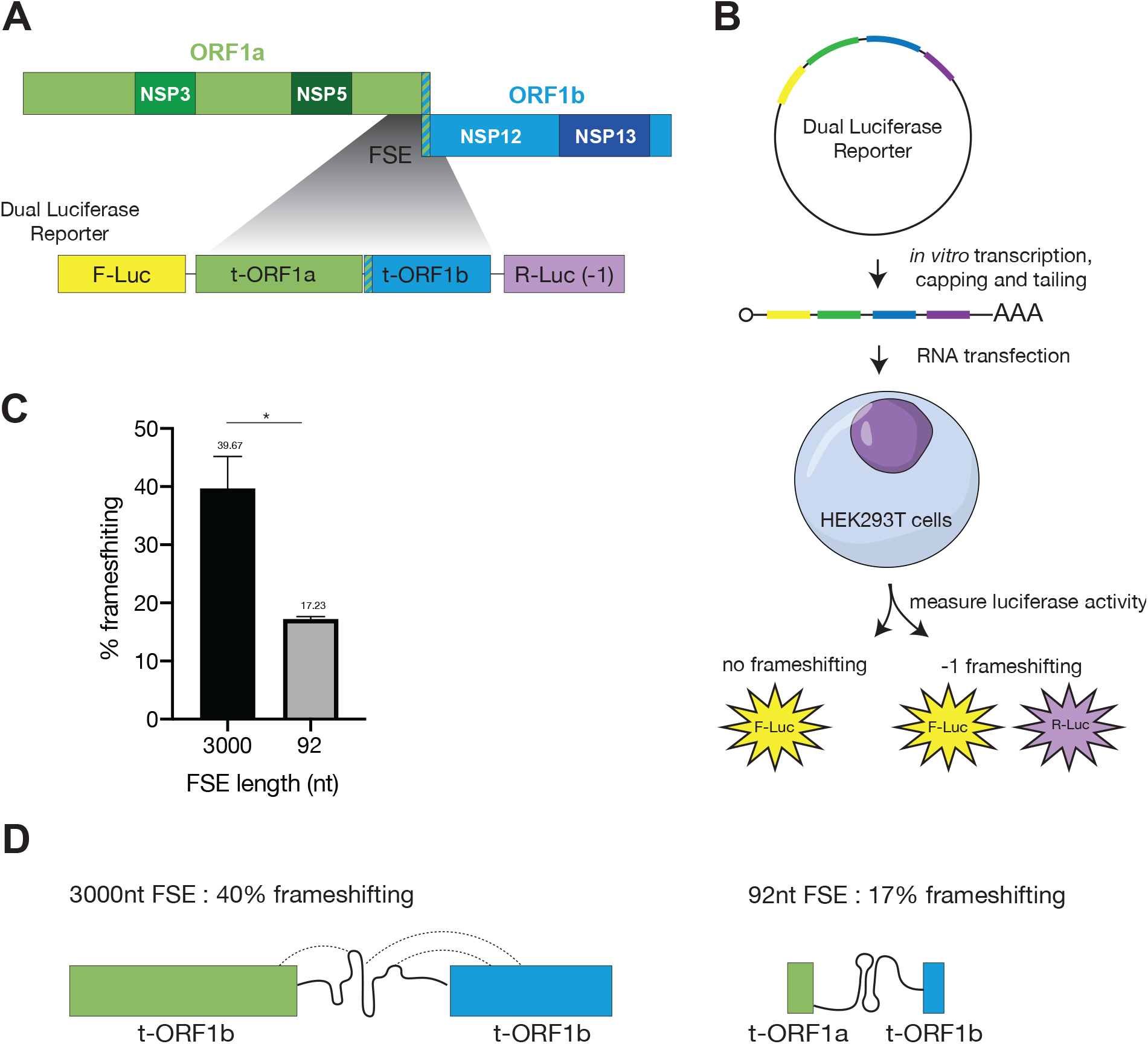
The long FSE element has dramatically higher frameshifting rate than the minimal FSE. **(A)** Schematic the long FSE. Truncated orf1ab (t-orf1ab) is inserted into a dual luciferase −1 frameshifting reporter. **(B)** The luciferase construct is in vitro transcribed, capped, tailed and transfected into HEK293T cells for 24h before measuring luciferase activity. No frameshifting results in only firefly luciferase (F-Luc) luminescence and −1 frameshifting results in F-Luc and Renilla luciferase (R-Luc) luminescence. **(C)** % frameshifting calculated as R-Luc/F-Luc % normalized against amino-acid matched positive control and negative control for both 92- and 3000nt FSE for n=3, p<0.05; *, unpaired t tests. **(D)** Schematic of RNA structure ensemble leading to higher frameshifting rate.

Previous studies using similar constructs have focused on just the short FSE and found that it promotes ~20% frameshifting (Kelly *et al.*, 2020; Sun *et al.*, 2020). Strikingly, we found that the long FSE frameshifted at ~40% while the short FSE frameshifted at only ~17% (Figure 6C). Our results on the long FSE are in agreement with *in vivo* ribosome profiling measurements of SARS-CoV-2 infected cells (Finkel *et al.*, 2021) (Figure 6C), indicating that the previously predicted structure of the canonical 92nt FSE does not recapitulate the mechanism of ribosomal frameshifting on the full-length virus during infection. Although additional studies are needed to understand the precise nature of the interactions between sequences further up and downstream in ORF1a and ORF1b that impact both the FSE structure ensemble and frameshifting rate (Figure 6D), our results underscore the importance of probing RNA secondary structure in cells and in its full-length context.

## Discussion

Here, we present the first insights into the secondary structure ensembles of the entire SARS-CoV-2 RNA genome in infected cells based on chemical probing with DMS-MaPseq. Previous work on the RNA structures of SARS-CoV-2 have provided only population-average models, which assume that the RNA folds into one conformation. In addition to our population-average model, we used the clustering algorithm DREEM (Tomezsko et al., 2020), and quantitatively detected alternative structures across the genome and revealed novel conformations at critical positions such as the frameshifting element (FSE).

Our DMS-MaPseq/DREEM framework gives the highest reproducibility data and agreement between the data and the predicted structure models, compared to all other chemical probing work on SARS-CoV-2 genome to date (Supplementary Figure 4). Importantly, our framework is the only approach that allows detecting RNA structure heterogeneity directly from the data itself, without any prior assumptions or thermodynamic and statistical modeling of RNA folding. We have previously benchmarked and validated DMS-MaPseq/DREEM on gold standard structures (Tomezsko *et al.*, 2020), and now we generate a secondary structure model for the entire SARS-CoV-2 genomic RNA, highlighting regions that are similar to well-folded RNAs as well as regions that are highly heterogeneous in folding.

Our in-cell data reveal alternative conformations for the frameshift element (FSE) within its genomic sequence context distinct from the canonical pseudoknot seen when considering only the 92nt FSE. We show that *in vitro* RNA-refolding of the full-length 30 kb genome can recapitulate the structure ensemble formed at the FSE in cells. Importantly, we show that the longer sequence is critical to achieve the frameshifting rate observed in cells during viral infection. When used in dual luciferase reporters, the longer sequence (3kb) frameshifts at much higher rate than the minimal FSE (~40% compared to ~20% of the minimal sequence). These results underscore a functional role for long range RNA interactions (Ziv *et al.*, 2020) and explain data from recent ribosome profiling studies showing that the ribosomes frameshifts at ~50% in infected cells (Puray-Chavez *et al.*, 2020; Finkel *et al.*, 2021).

Our in-cell data-derived model of SARS-CoV-2 presents major RNA structures and sites of RNA structure heterogeneity across the entire genome and provides the foundation for further studies. Importantly, our work reveals that drugs such as small molecules or anti-sense oligoes intended to abolish SARS-CoV-2 frameshifting should be designed and tested against the correct structure ensemble that forms in cells. Further work to better understand of the functional significance of other structured elements across SARS-CoV-2 genome will enable the design of more targeted therapeutics.

## Acknowledgments

The SARS-CoV-2 starting material was provided by the World Reference Center for Emerging Viruses and Arboviruses (WRCEVA), with Natalie Thornburg as the CDC Principal Investigator.” We thank T.B. Faust for manuscript input and E. Smith for illustrator images. This work was supported by The Pershing Square Foundation, The Office of Naval Research Award # N00014-20-1-2084, and the Burroughs Wellcome fund.

## Author Contributions

T.C.T.L. and S.R. conceived and designed the project. T.C.T.L. carried out all experiments with collaborative contributions from L.E.M. and A.G. M.F.A., T.C.T.L., S.K., S.S.Y.N. and S.R. performed the data analysis. J.G. and Y.S. performed the reporter frameshifting assays. T.C.T.L., M.F.A., and S.R. interpreted the results and wrote the paper with input from S.K., S.S.Y.N., L.E.M, M.B., and A.G.

## Methods

### RESOURCE AVAILABILITY

#### Lead Contact

Further information and requests for resources and reagents should be directed to and will be fulfilled by the Lead Contact, S. Rouskin

#### Materials Availability

This study did not generate new unique reagents.

#### Data and Code Availability

The source code for the data processing and analyses is available at http://dreem.wi.mit.edu/static/dreem.zip and http://dreem.wi.mit.edu/static/DREEM_Manual.pdf

The sequencing data are deposited into NCBI Gene Expression Omnibus (GEO), (accession number pending).

### EXPERIMENTAL MODEL AND SUBJECT DETAILS

SARS-CoV-2 total viral RNA was extracted from Vero cells (ATCC CCL-81) cultured in DMEM (Gibco) supplemented with 10% FBS (Gibco) plated into 100 mm dishes and infected at a MOI of 0.01 with 2019-nCoV/USA-WA1/2020 (Passage 6). Infected cells were incubated at 37 °C, 5% CO_2_ and harvested 2 days post infection either with or without DMS treatment. Infected cell pellets were centrifuged at 5000xg for 5 min at 4 °C and resuspended in Trizol (Ambion).

### METHOD DETAILS

#### DMS modification of SARS-CoV-2 RNA in infected cells

200 μl DMS (or 2% v/v) was added dropwise to the plated Vero cells 48 h post SARS-CoV-2 infection and incubated for 4 min at 37°C. DMS was neutralized by adding 15 ml PBS (ThermoFisher Scientific) with 30% β-mercaptoethanol. The cells were centrifuged at 1,000g for 5 min at 4°C. The cells were washed twice by resuspending the pellet with 15 ml PBS with 30% β-mercaptoethanol and centrifugation to pellet then just once with 15 ml PBS. After washes, the pellet was resuspended in 1 ml Trizol (ThermoFisher Scientific) and RNA was extracted following the manufacturer’s specifications. Total RNA was purified using RNA Clean and Concentrator −25 kit (Zymo).

#### DMS modification of *in vitro*-transcribed RNA

gBlocks were obtained from IDT for the SARS-CoV-2 92nt and 283nt FSE which corresponds to nucleotides 13460-13546 and nucleotides 13,342-13,624 based on 2019-nCoV/USA-WA1/2020. The regions of interest were amplified by PCR with a forward primer that contained the T7 promoter sequence (TAATACGACTCACTATAGGGTT). The PCR product was used for T7 Megascript in vitro transcription (ThermoFisher Scientific) according to manufacturer’s instructions with a 16 h incubation time at 37 °C. Subsequently, 1 μl Turbo DNase I (ThermoFisher Scientific) was added to the reaction and incubated at 37°C for 15 min. The RNA was purified using RNA Clean and Concentrator −5 kit (Zymo). 10 μg of RNA in 10 μl H_2_O was denatured at 95°C for 1 min then placed on ice. On the basis of the DMS concentration used in the next step, 300 mM sodium cacodylate buffer (Electron Microscopy Sciences) with 6 mM MgCl_2_+ (refolding buffer) was added so that the final volume was 100 μl. (e.g. for 2.5% final DMS concentration: add 87.5 μl refolding buffer and 2.5 μl DMS) Then, 2.5 μl was added and incubated at 37°C for 5 min while shaking at 500 r.p.m. on a thermomixer. The DMS was neutralized by adding 60 μl β-mercaptoethanol (Millipore-Sigma). The RNA was purified using RNA Clean and Concentrator −5 kit.

#### DMS modification of full-length SARS-CoV-2 RNA *in vitro*

Full-length SARS-CoV-2 RNA was extracted from the supernatant of infected Vero cells (as described above), resuspended in 1 ml Trizol (ThermoFisher Scientific) and RNA was extracted following the manufacturer’s specifications. The RNA was purified using RNA Clean and Concentrator −5 kit (Zymo) and DMS modified as described above.

#### Human rRNA subtraction of total cellular RNA

15 μg of total RNA per reaction was used as the input for rRNA subtraction. First, 1 μl rRNA subtraction mix (15 μg/μl) and 2 μl 5× hybridization buffer (end concentration: 200 mM NaCl, 100 mM Tris-HCl, pH 7.4) were added to each reaction, and final volume was then adjusted with water to 10 μl. The samples were denatured at 95°C for 2 min and then temperature was reduced by 0.1°C/s until the reaction was at 45°C. Next, 10 μl RNase H buffer and 2 μl hybridase thermostable RNase H (Lucigen) preheated to 45° were added. The samples were incubated at 45°C for 30 min. The RNA was cleaned with RNA Clean and Concentrator −5, following the manufacturer’s instructions and eluted in 45 μl water. Then, 5 μl Turbo DNase buffer and 3 μl Turbo DNase (ThermoFisher Scientific) were added to each reaction and incubated for 30 min at 37°C. The RNA was purified with RNA Clean and Concentrator −5 (Zymo) following instructions.

#### RT–PCR and sequencing of DMS-modified RNA

For reverse transcription, 1.5 μg of rRNA subtracted total RNA or 10 μg of in vitro-transcribed RNA was added to 4 μl 5× first strand buffer (ThermoFisher Scientific), 1 μl 10μM reverse primer, 1 μl dNTP, 1 μl 0.1M DTT, 1 μl RNaseOUT and 1 μl TGIRT-III (Ingex). The reverse-transcription reaction was incubated at 60°C for 1.5 h. 1 μl 4M NaOH was then added and incubated at 95°C for 3 min to degrade the RNA. The cDNA was purified with Oligo Clean and Concentrator −5 (Zymo) following instructions. PCR amplification was done using Advantage HF 2 DNA polymerase (Takara) for 30 cycles according to the manufacturer’s specifications. The PCR product was purified by DNA Clean and Concentrator −5 (Zymo) following manufacturer’s instructions. RNA-seq library for 150 bp insert size was constructed following the manufacturer’s instruction (NEBNext Ultra™ II DNA Library Prep Kit). The library was loaded on ISEQ-100 Sequencing flow cell with ISEQ-100 High-throughput Sequencing Kit and the library was run on ISEQ-100 (paired-end run,151 × 151 cycles).

#### Library generation with DMS-modified SARS-CoV-2 RNA

After rRNA subtraction (described above), extracted DMS-modified RNA from SARS-CoV-2 infected Vero cells was fragmented using the RNA Fragmentation kit (ThermoFisher Scientific). 1.5 μg of rRNA subtracted total RNA was fragmented at 70°C for 2.5 min. The fragmented RNA was mixed with an equal volume 2× Novex TBE-urea sample buffer (ThermoFisher Scientific) and run on a 10% TBE-urea gel (ThermoFisher Scientific) at 200V for 1 h 15 min for size selection of RNA that is ~150nt. To dephosphorylate and repair the ends of randomly fragmented RNA, 2 μl 10x CutSmart buffer (New England Biolabs), 10 μl shrimp alkaline phosphatase (New England Biolabs), 2 μl RNaseOUT (ThermoFisher Scientific) and water were added to a final volume of 20 μl and 37°C for 1 h. Next, 4 μl 50% PEG-800 (New England Biolabs), 4 μl 10× T4 RNA ligase buffer (New England Biolabs), 4 μl T4 RNA ligase, truncated KQ (England Biolabs) and 2 μl linker were added to the reaction and incubated for 18 h at 22°C. The RNA was purified with RNA Clean and Concentrator −5, following the manufacturer’s instructions for recovery of all fragments and eluted in 10 μl water. Excess linker was degraded by adding 2 μl 10× RecJ buffer (Lucigen), 1 μl RecJ exonuclease (Lucigen), 1 μl 5′ deadenylase (New England Biolabs) and 1 μl RNaseOUT, then incubating for 1 h at 30°C. The RNA was purified with RNA Clean and Concentrator −5, following the manufacturer’s instructions and eluted in 11 μl water.

For reverse transcription, 1.5 μg of rRNA subtracted total RNA or 10 μg of in vitro-transcribed RNA was added to 4 μl 5× first strand buffer (ThermoFisher Scientific), 1 μl 10μM reverse primer, 1 μl dNTP, 1 μl 0.1M DTT, 1 μl RNaseOUT and 1 μl TGIRT-III (Ingex). The reverse-transcription reaction was incubated at 60°C for 1.5 h. 1 μl 4M NaOH was then added and incubated at 95°C for 3 min to degrade the RNA. The reverse-transcription product was mixed with an equal volume 2× Novex TBE-urea sample buffer (ThermoFisher Scientific) and run on a 10% TBE-urea gel (ThermoFisher Scientific) at 200V for 1 h 15 min for size selection of cDNA that is ~250nt. The size-selected and purified cDNA was circularized using CircLigase ssDNA ligase kit (Lucigen) following manufacture’s protocol. 2 μl of the circularized product was then used for PCR amplification using Phusion High-Fidelity DNA Polymerase (NEB) for a maximum of 16 cycles. The PCR product was run on an 8% TBE gel at 180V for 1 h and size-selected for products ~300 nt. The product was then sequenced with iSeq100 (Illumina) to produce either 150×150-nt paired-end reads.

#### Dual-luciferase frameshift reporter assay

92nt and 3000nt FSEs which corresponds to nucleotides13460-13546 and nucleotides 12686-15609 based on 2019-nCoV/USA-WA1/2020 were inserted into dual luciferase reporter bewteen firefly luciferease (F-Luc) coding sequence and renilla luciferase (R-Luc) coding sequence in −1 frame. Insertion of 0-frame stop codon between FLuc and FSE element is used as negative control construct whilst a construct of matching length in which F-Luc and R-Luc were translated continuously without frameshifting is used as a positive control.

Frameshifting reporter as well as positive and negative control mRNAs were in vitro transcribed and polyadenylated using HiScribe T7 mRNA kit (New England Biolabs) according to the manufacturers’ instructions. Purified mRNAs were transfected in HEK293T cells in 24-well plates using Lipofectamine MessengerMAX (ThermoFisher). 24 hours after transfection, cells were washed once with phosphate-buffered saline (PBS), and lysed in Glo Lysis Buffer (Promega) at room temperature for 5 min. 10 μL of lysate was diluted with 30 μL PBS before being mixed with 40 μL Dual-Glo FLuc substrate (Promega). After 10 min, FLuc activity was measured in a GloMax 20/20 luminometer (Promega). Subsequently, 40 μL Dual-Glo Stop & Glo reagent was added to the mixture, incubated for 10 min, and measured for RLuc luminescence. The ratio between RLuc and FLuc activities minus the negative control background luminescence and normalized to positive control luminescence was calculated as frameshift efficiency.

### QUANTIFICATION AND STATISTICAL ANALYSIS

#### Mapping and quantification of mutations

Fastq files were trimmed using TrimGalore (github.com/FelixKrueger/TrimGalore) to remove Illumina adapters. Trimmed paired reads were mapped to the genome of SARS-CoV-2 isolate SARS-CoV-2/human/USA/USA-WA1/2020 (GenBank: MN985325.1) (Harcourt *et al.*, 2020) using Bowtie2 (Langmead and Salzberg, 2012) with the following parameters: --local --no-unal --no-discordant --no-mixed -L 12 -X 1000. Reads aligning equally well to more than one location were discarded. SAM files from Bowtie2 were converted into BAM files using Picard Tools SamFormatConverter (broadinstitute.github.io/picard).

For each pair of aligned reads, a bit vector the length of the reference sequence was generated using DREEM (Tomezsko *et al.*, 2020). Bit vectors contained a 0 at every position in the reference sequence where the reference sequence matched the read, a 1 at every base at which there was a mismatch or deletion in the read, and no information for every base that was either not in the read or had a Phred score <20. We refer to positions in a bit vector with a 0 or 1 as “informative bits” and all other positions as “uninformative bits.”

For each position in the reference sequence, the number of bit vectors covering the position and the number of reads with mismatches and deletions at the position were counted using DREEM. The ratio of mismatches plus deletions to total coverage at each position was calculated to obtain the population average mutation rate for each position.

#### Filtering bit vectors

In cases indicated below, bit vectors were discarded if they had two mutations closer than 4 bases apart, had a mutation next to an uninformative bit, or had more than an allowed total number of mutations (greater than 10% of the length of the bit vector and greater than three standard deviations above the median number of mutations among all bit vectors). The average mutation rate for each position was computed from the filtered bit vectors in the same way as described above.

#### Normalizing the mutation rates

The mutation rates for all of the bases in the RNA molecule were sorted in numerical order. The greatest 5% or 10% of mutation rates (specified where relevant in the main text) were chosen for normalization. The median among these signals was calculated. All mutation rates were divided by this median to compute the normalized mutation rates. Normalized rates greater than 1.0 were winsorized by setting them to 1.0 (Dixon, 1960).

#### Computing genome coverage and mutation rates

Genome-wide coverage (Figure 1C) was computed by counting the number of unfiltered bit vectors from the in-cell library that contained an informative bit (0 or 1) at each position. Signal and noise plots (Figure 1D) were generated from the unfiltered population average mutation rate. A total of 103 (0.34%) positions across the genome were discarded for having a noise mutation rate greater than 1% in the untreated sample (likely due to endogenous modifications or “hotspot” reverse transcription errors). The signal and noise were computed every 100 nt, starting at nucleotide 51. For each of these nucleotides, the average mutation rate was computed over the 100 nt window starting 50 bases upstream and ending 49 bases downstream. The “signal” was defined as the average mutation rate of A and C, while the “noise” was defined as the average mutation rate of G and U.

The correlation of mutation rates between biological replicates genome-wide (Figure 1B) was computed using the unfiltered bit vectors. The correlation of mutation rates between different conditions of the FSE (Figure 4B) was computed using the filtered bit vectors. The correlation of mutation rates between clusters and biological replicates for the FSE (Figure 5B) was computed using the filtered bit vectors after clustering into two clusters. For all correlation plots, the Pearson correlation coefficient is given. A total of 6 (0.02%) outliers with >30% mutation rate were removed to prevent inflating the Pearson correlation coefficients.

#### Folding the entire SARS-CoV-2 genome

The unfiltered population average mutation rate was obtained from the in-cell library reads. The 29,882 nt genome of SARS-CoV-2 was divided into ten segments, each roughly 3 kb the boundaries of which are predicted to be open and accessible by RNAz (Rangan, Zheludev and Das, 2020). For each segment, the population average mutation rate was normalized. The segment was then folded using the Fold algorithm from RNAstructure (Mathews, 2004) with parameters -m 3 to generate the top three structures, -md to specify a maximum base pair distance, and -dms to use the normalized mutation rates as constraints in folding. All mutation rates on G and U bases were set to −999 (unavailable constrains). Connectivity Table files output from Fold were converted to dot bracket format using ct2dot from RNAstructure (Mathews, 2004). The ten dot bracket structures were concatenated into a single genome-wide structure.

#### The data-structure correlation index (DSCI)

The data-structure correlation index (DSCI) quantifies how well a secondary structure model is supported by DMS or SHAPE reactivity data, under the assumption that genuinely unpaired bases are more reactive than paired bases. Given a secondary structure model in which every base is designated as paired or unpaired, and reactivity values for all or for a subset of bases in the model, the DSCI is defined as the probability that a randomly chosen unpaired base will have greater reactivity than a randomly chosen paired base. It is equal to the following:

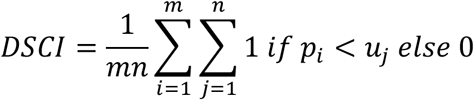

where *p* is the set of reactivities for all *m* paired bases (indexed by *i*) and *u* is the set of reactivities all *n* paired bases (indexed by *j*). Bases without reactivity information (such as Gs and Us for DMS data, and any problematic base) are excluded from *p* and *u*.

The DSCI is closely related to the Mann-Whitney *U* statistic (Mann and Whitney, 1947), which is obtained from the above equation without dividing by *mn* (assuming no ties in reactivities). The calculation is implemented in Python using the SciPy Stats MannWhitneyU function (Virtanen *et al.*, 2020), and dividing the result by *mn*. If min(*m*, *n*) < 5, then we return a missing value to avoid biases caused by very low numbers of paired or unpaired bases.

#### The modified Fowlkes-Mallows index (mFMI)

Given two RNA structures of the same length (*L*) in dot-bracket notation, all base pairs in each structure were identified. Each base pair was represented as a tuple of (position of 5’ base, position of 3’ base). The number of base pairs common to both structures (*P_12_*) as well as the number of base pairs unique to the first structure (*P_1_*) and to the second structure (*P_2_*) were computed. Given these quantities, the Fowlkes-Mallows index (a measure of similarity between two binary classifiers) is defined as 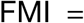 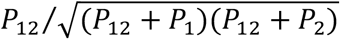 (Fowlkes and Mallows, 1983). In the case that (*P*_12_ + *P*_1_)(*P*_12_ + *P*_2_) = 0, we let FMI = 0.

As the Fowlkes-Mallows index does not consider positions at which the structures agree on bases that are unpaired, the index needed to be modified; otherwise regions with few base pairs would tend to score too low. Thus, the number of positions at which both sequences contained an unpaired base (*U*) was computed. Two variations of the modified Fowlkes-Mallows index (mFMI) were tested that differed in their treatment of externally paired bases, defined as bases paired to another base outside of the region of the structure being compared. The version of mFMI excluding external base pairs counted all externally paired bases as unpaired when computing *U*. The number of positions containing a paired base (*P*) was computed as *P* = *L* – *U*. In this case, mFMI was defined as mFMI = *U*/*L* + *P*/*L* × FMI, which weights the Fowlkes-Mallows index by the fraction of paired bases and adds the fraction of unpaired bases (*U*/*L*), as the structures agree at all unpaired positions.

To include external base pairs, any position containing an externally paired base was not counted in *U*. The number of positions at which both structures contained an externally paired base with the same orientation (i.e. both facing in the 5’ or 3’ direction) was computed as the number *E*. The number of positions at which at least one structure contained a base that was paired, but not externally, was computed as *P*. Then, the mFMI was defined as mFMI = *U*/*L* + *E*/*L* + *P*/*L* × FMI, which weights the Fowlkes-Mallows index by the fraction of positions containing a paired base and considers positions in which both bases are unpaired as in agreement, but only counts externally paired bases as agreeing if both structures contain an externally paired base at the same position and the base pairs have the same orientation.

#### Comparisons to previous *in silico* predictions

Excel files from the supplemental material of (Rangan, Zheludev and Das, 2020) were parsed to obtain the coordinates and predicted structures. For each predicted structure, agreement with the region of our structure with the same coordinates was computed using the mFMI, either including or excluding external base pairs (as specified in the text). Box plots of the agreement for each window (Figure 3B) show the minimum, first quartile, median, third quartile, and maximum; data lying more than 1.5 times the interquartile range from the nearest quartile are considered outliers and are plotted as individual points. The numbers of points in each box plot are given in the Results section for Figure 3B.

#### Folding the frameshift stimulating element

Reads from RT-PCR of a 283 nt segment of in-cell RNA spanning the FSE (nucleotides 13,342 - 13,624) were used to generate bit vectors. The bit vectors were filtered as described above, and the filtered average mutation rates were normalized. The RNA was folded using the ShapeKnots algorithm from RNAstructure (Hajdin *et al.*, 2013) with parameters -m 3 to generate three structures and -dms to use the normalized mutation rates as constraints in folding. All signals on G and U bases were set to −999 (unavailable constrains). Connectivity Table files output from Fold were converted to dot bracket format using ct2dot from RNAstructure (Mathews, 2004).

#### Coronavirus sequence alignments

Accession numbers of curated sarbecovirus and merbecovrus genomes were obtained from (Ceraolo and Giorgi, 2020) and downloaded from NCBI. The sequences were aligned using the MUSCLE (Edgar, 2004) web service with default parameters. The region of the multiple sequence alignment spanning the two sides of Alternative Stem 1 was located and the sequence conservation computed using custom Python scripts.

For the alignment of all betacoronaviruses with genomes in NCBI RefSeq (O’Leary *et al.*, 2016), all reference genomes of betacoronaviruses were downloaded from RefSeq using the query “betacoronavirus[organism] AND complete genome” with the RefSeq source database as a filter. The sequences were aligned using the MUSCLE (Edgar, 2004) web service with default parameters. The subgenus of betacoronavirus to which each virus belonged was obtained from the NCBI taxonomy database (Sayers *et al.*, 2009).

#### Detecting alternative structures genome-wide

The reference genome (length = 29,882 nt) was partitioned into 373 regions of 80 nt each and one final region of 42 nt. For each region, reads were filtered out according to the criteria in “Filtering Bit Vectors” or if they did not overlap with at least 20% (16 nt) of the region. The reads were then clustered using the EM algorithm implemented previously (Tomezsko *et al.*, 2020) using a maximum of two clusters per region, ignoring G and U residues, and setting all mutation rates less than 0.005 to 0.0.

After clustering, regions were filtered out if fewer than 100,000 reads mapped to the region (n = 42) or if either cluster contained a base with a mutation rate exceeding 30% (n = 16). For each remaining region with two clusters (n = 316), each cluster’s mutation rates (μ) were normalized by setting the base with the highest mutation rate to 1.0 and scaling the mutation rates of all other bases proportionally. For each base, the difference in DMS reactivities (ΔDMS) between its mutation rate in cluster 1 (μ_1_) and cluster 2 (μ_2_) was calculated as 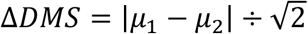. The coefficient of determination (R^2^) was also computed on the normalized DMS reactivities.

#### Detecting alternative structures of the FSE

The filtered bit vectors (the same used to fold the frameshift stimulating element) were clustered using the expectation maximization algorithm of DREEM to allow detection of a maximum of two alternative structures (Tomezsko *et al.*, 2020).

#### Quantification of minus-strand reads

Mapped reads from the in-cell library were classified as minus-strand using a custom Python script if they had the following SAM flags (Li *et al.*, 2009): PAIRED and PROPER_PAIR and ({READ1 and MREVERSE and not REVERSE} or {READ2 and REVERSE and not MREVERSE}) and not (UNMAP or MUNMAP or SECONDARY or QCFAIL or DUP or SUPPLEMENTARY).

#### Visualizing RNA structures

RNA structures were drawn using VARNA (Darty, Denise and Ponty, 2009). The bases were colored using the normalized DMS signals.

## Supplementary Figure legends

**Supplementary Figure 1: In-cell data-derived secondary structure of the full SARS-CoV-2 genome.**

**Supplementary Figure 2: Genome-wide data-structure correlation index (DSCI) for population average models from this study, Huston et al., and Manfredonia et al.**

**Supplementary Figure 3: Genome-wide pairwise similarity of population average models from this study, Huston et al., and Manfredonia et al.**

**Supplementary Figure 4: Comparison of our in-cell genome-wide structure model with previous computational models**

**(A)** Consistency of our in-cell structure models. Agreement is given between our structure models predicted using a maximum distance limit of 120 nt and 350 nt between paired bases at 5% signal normalization and between our predictions using 5% and 10% DMS normalization at 350 nt maximum allowed base pair distance.

**(B)** Agreement of our structure model with all predicted structures from RNAz and Contrafold. Agreement is given for both excluding and including external base pairs.

**(C)** Agreement of our structure with a previous model from RNAz across the genome. At positions for which multiple RNAz model exists, the average agreement with all models is given.

**(D)** Agreement of our model with RNAz predicted structures with the three highest P-values in regions with previously unannotated structures.

**(E)** Agreement of our model with Contrafold predicted structures with the five highest maximum expected accuracies in evolutionarily conserved regions.

**(F)** Agreement of our TRS structure models to RNAz predicted structures. For TRSs for which multiple RNAz models exist, agreement with each prediction is shown.

**Supplementary Figure 5: Structured and unstructured regions in the SARS-CoV-2 genome.**

**(A)** Locations of highly structured and unstructured regions in the SARS-CoV-2 genome. Highly structured regions are defined are stretches of at least 10 consecutive paired bases; unstructured regions shown are stretches of at least 14 consecutive unpaired bases. The thickness of each bar is proportional to the number of consecutive paired (blue) or unpaired (orange) bases. The data is plotted over a schematic of the genome, highlighting the organization of open reading frames (ORFs) and the transcription regulatory sequences (TRS).

**(B)** In-cell model of each of the eight TRSs predicted to lie within a stem loop. The core sequence (CS) of each TRS is outlined in black. Models are arranged in genomic order from top-to-bottom, left-to-right.

**Supplementary Figure 6: Relationship of DSCI and median ΔDMS for every overlapping 80nt window genome-wide**

