## Supplementary Figure 1 for "Insights into the secondary structural ensembles of the full SARS-CoV-2 RNA genome in infected cells"

### Supplementary Figure legends

**Supplementary Figure 1: In-cell data-derived secondary structure of the full SARS-CoV-2 genome.**

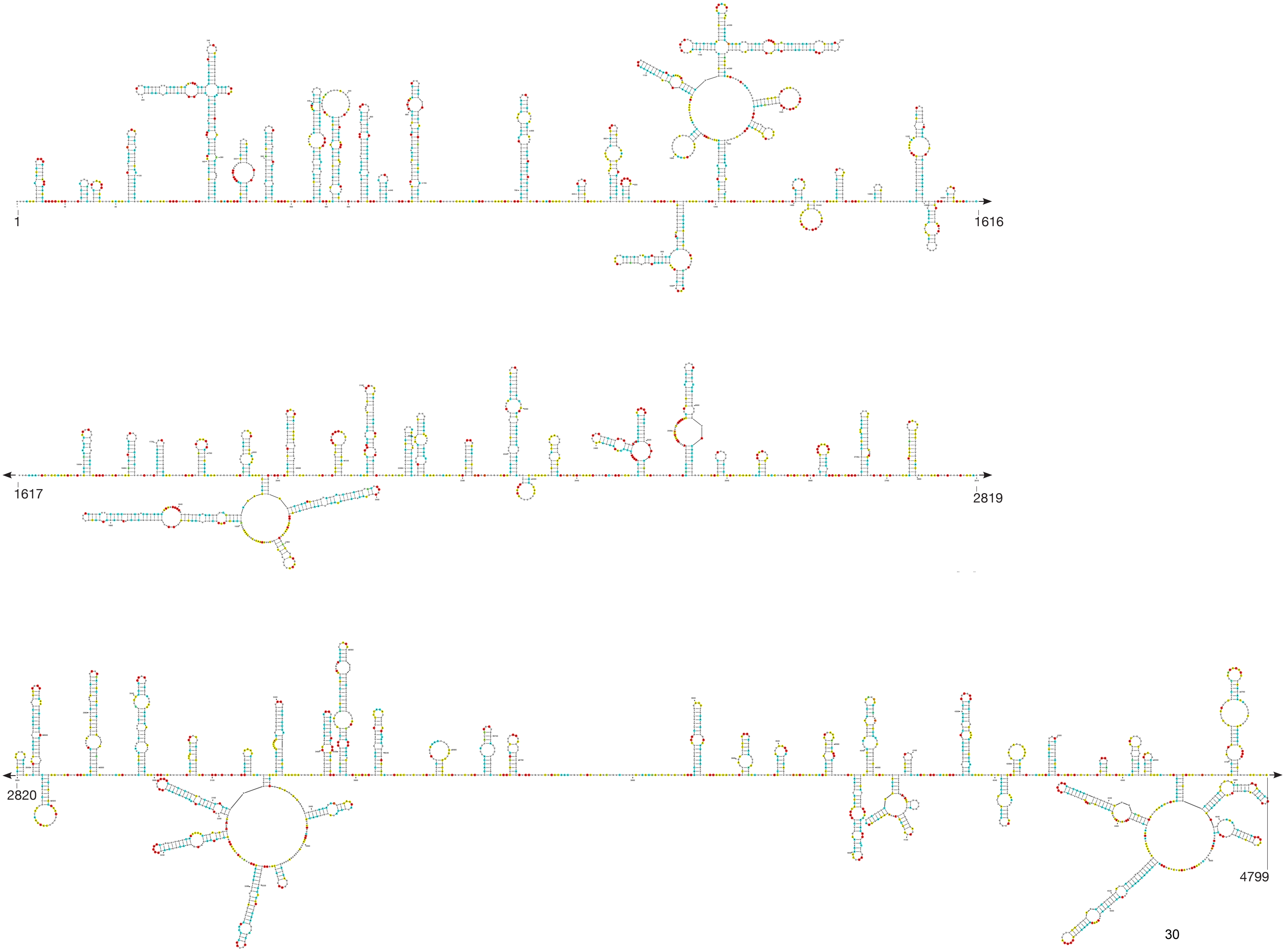
1
1616
1617
2819
2820
4799

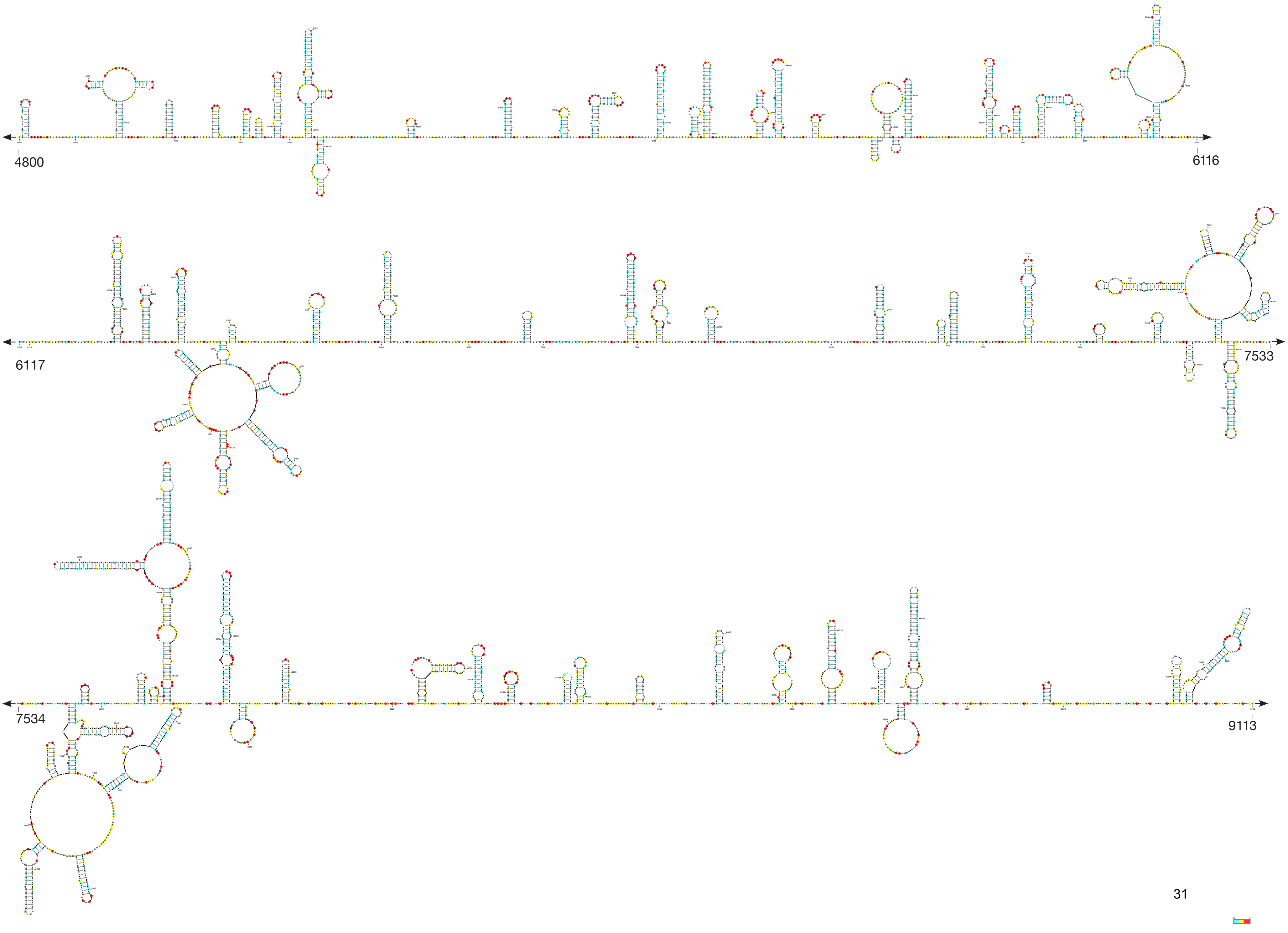

4800
6116
6117
7533
7534
9113

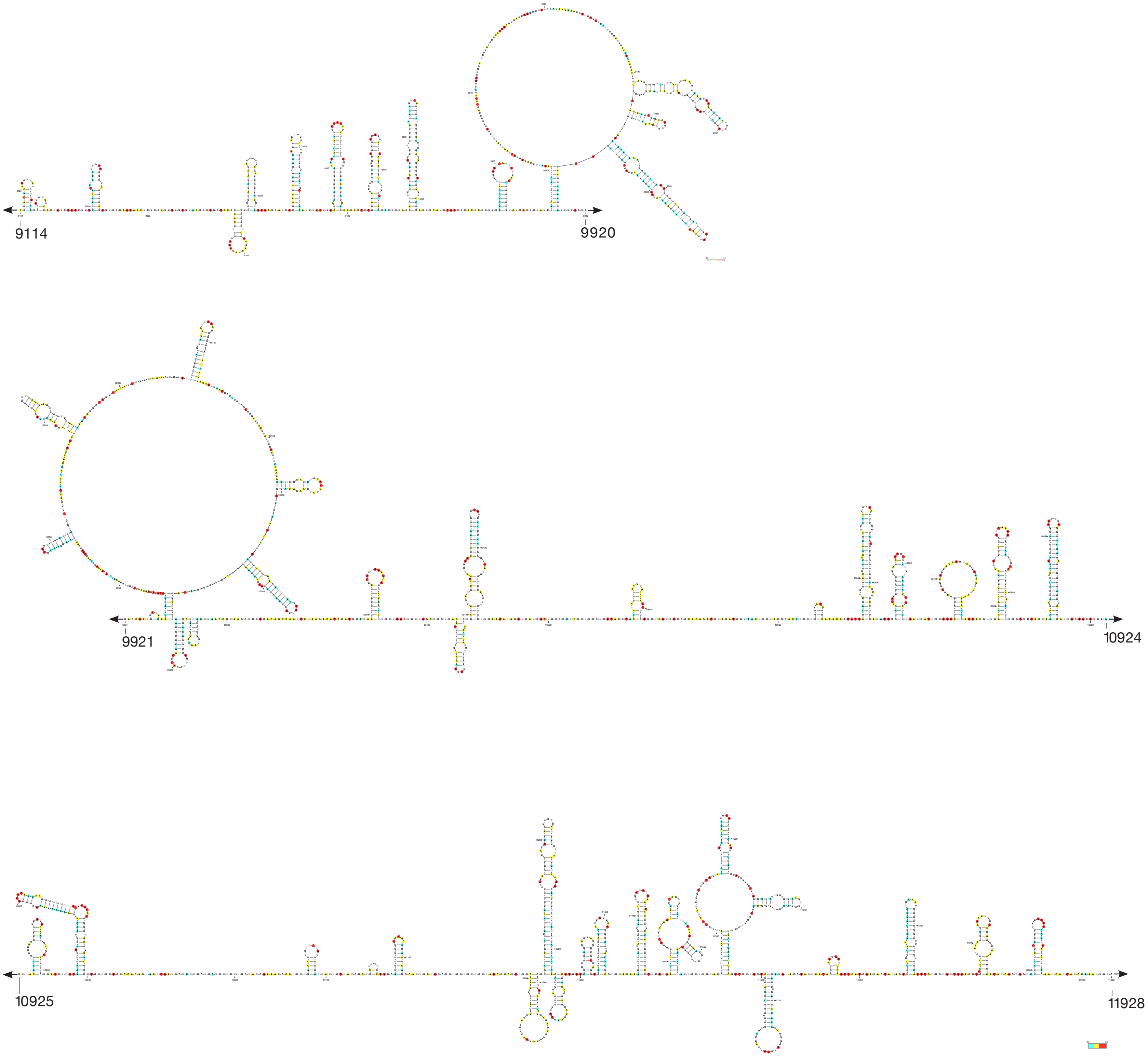
9114
9920
9921
10924
10925
11928

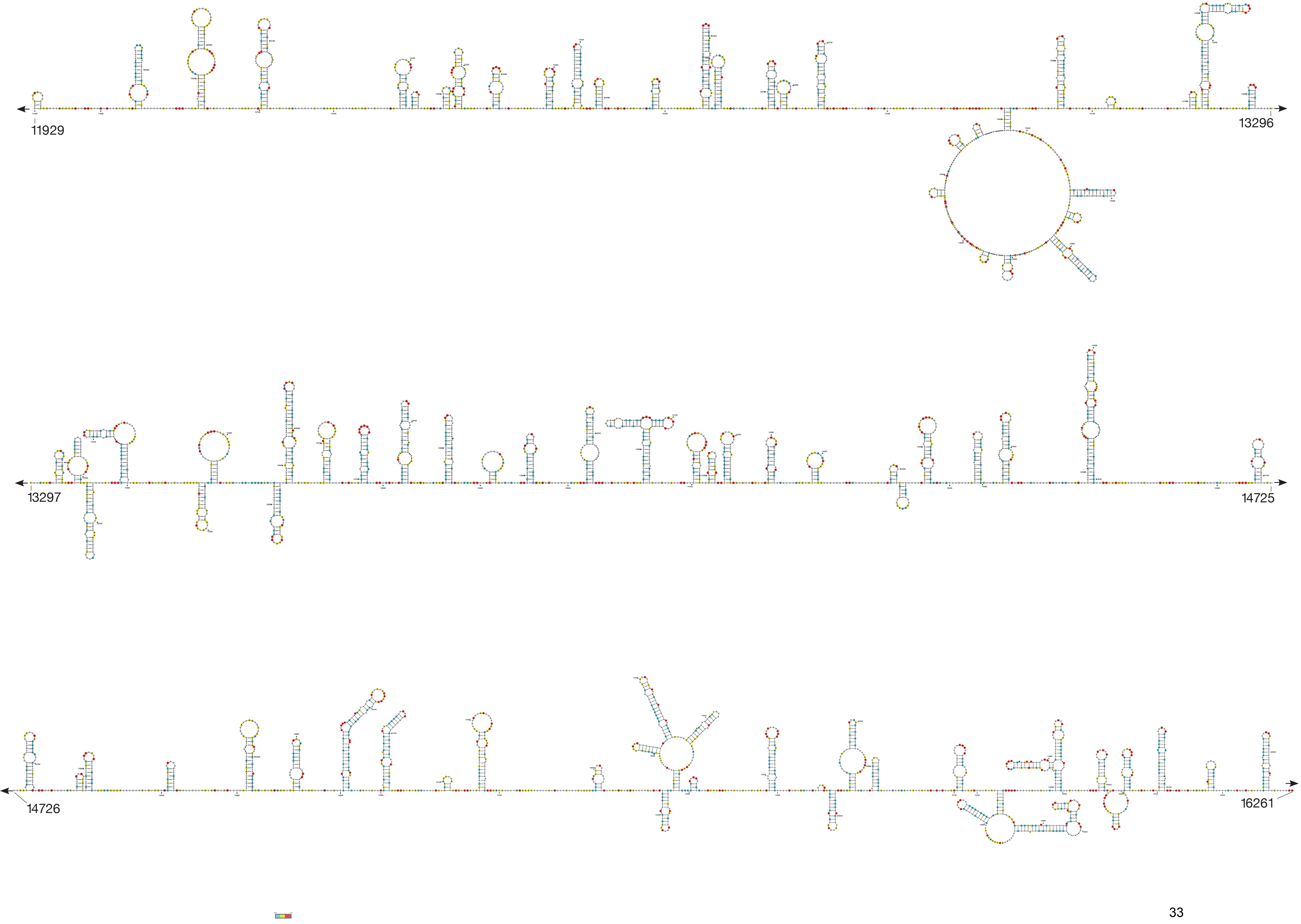

11929
13296
13297
14725
14726
16261

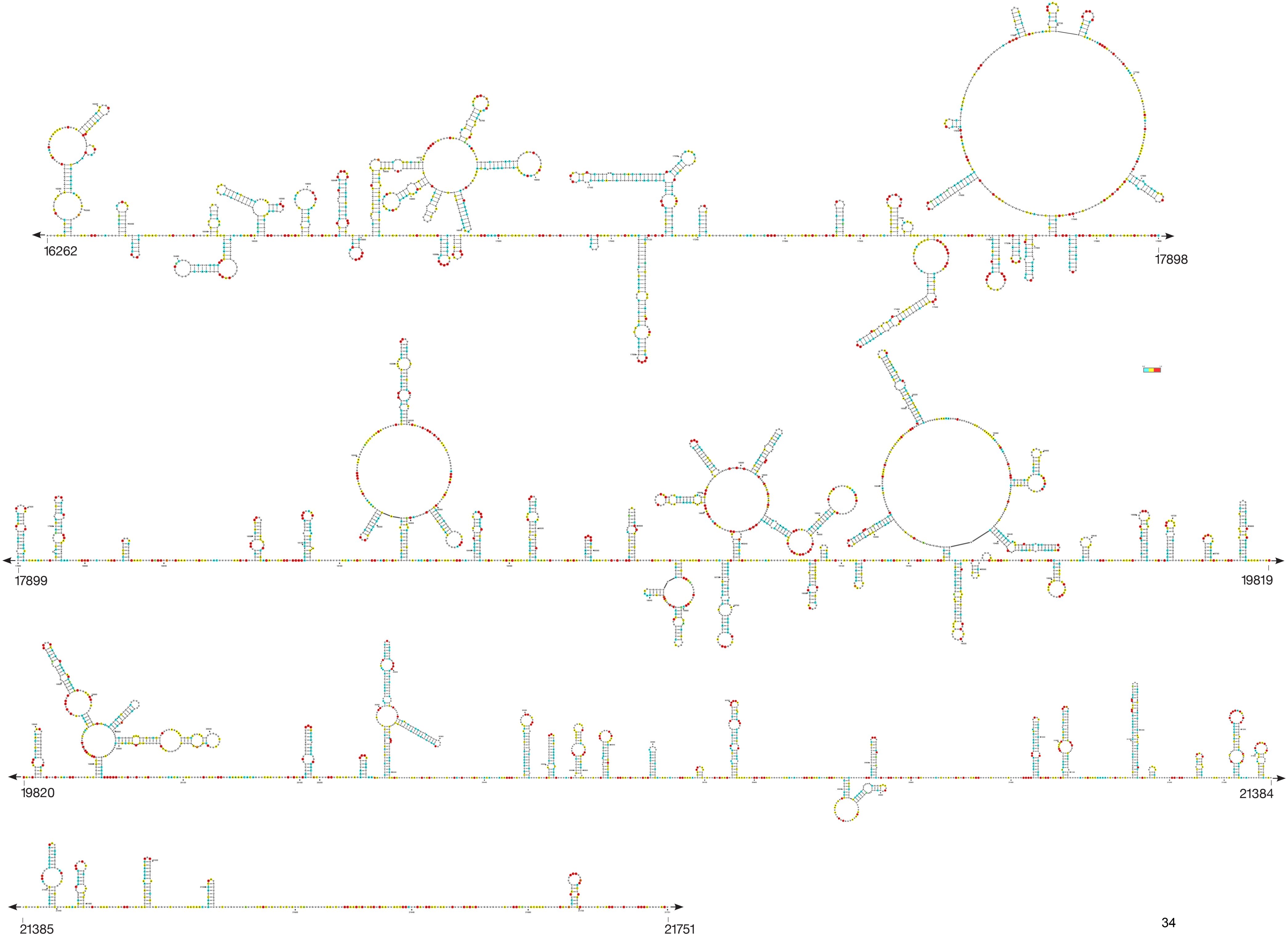
16262
17898
17899
19819
19820
21384
21385
21751

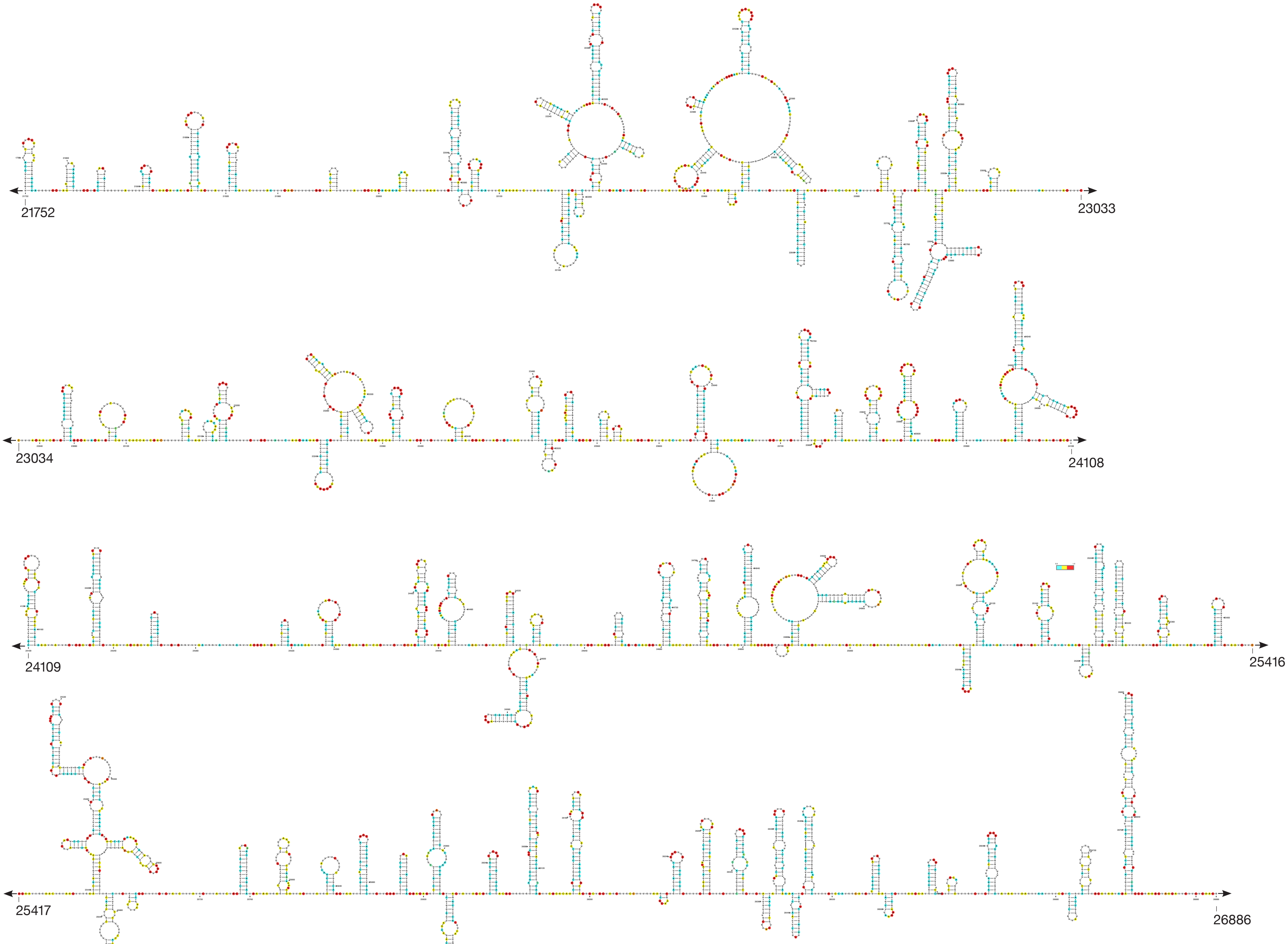


35

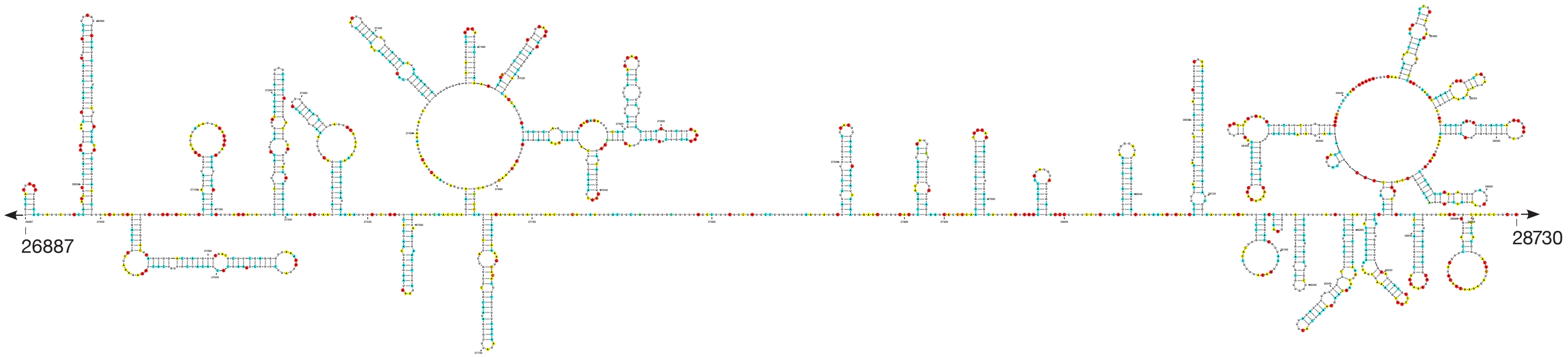
26887
28730

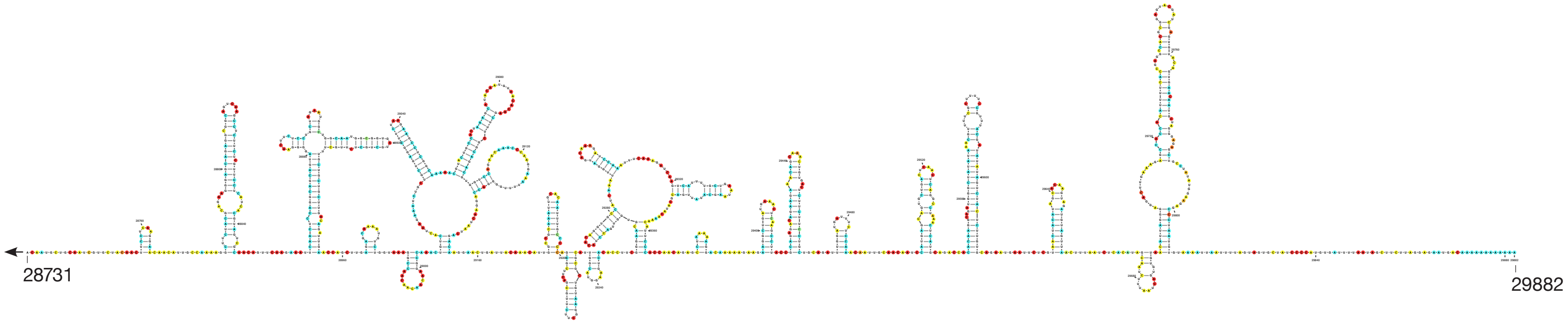
28731
29882
