## Supplementary Figures 2 - 6 for "Insights into the secondary structural ensembles of the full SARS-CoV-2 RNA genome in infected cells"

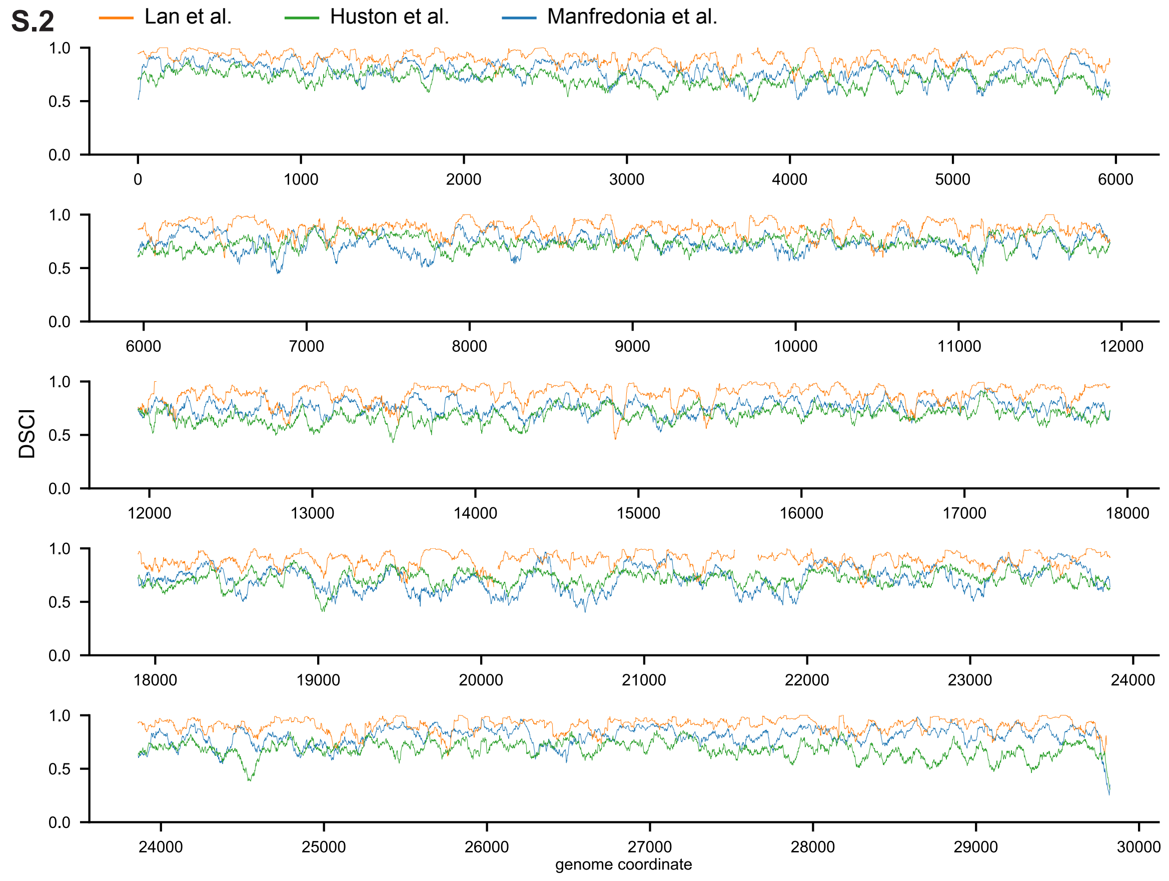

S.2
Lan et al.
Huston et al.
Manfredonia et al.
DSCI
genome coordinate

S.3

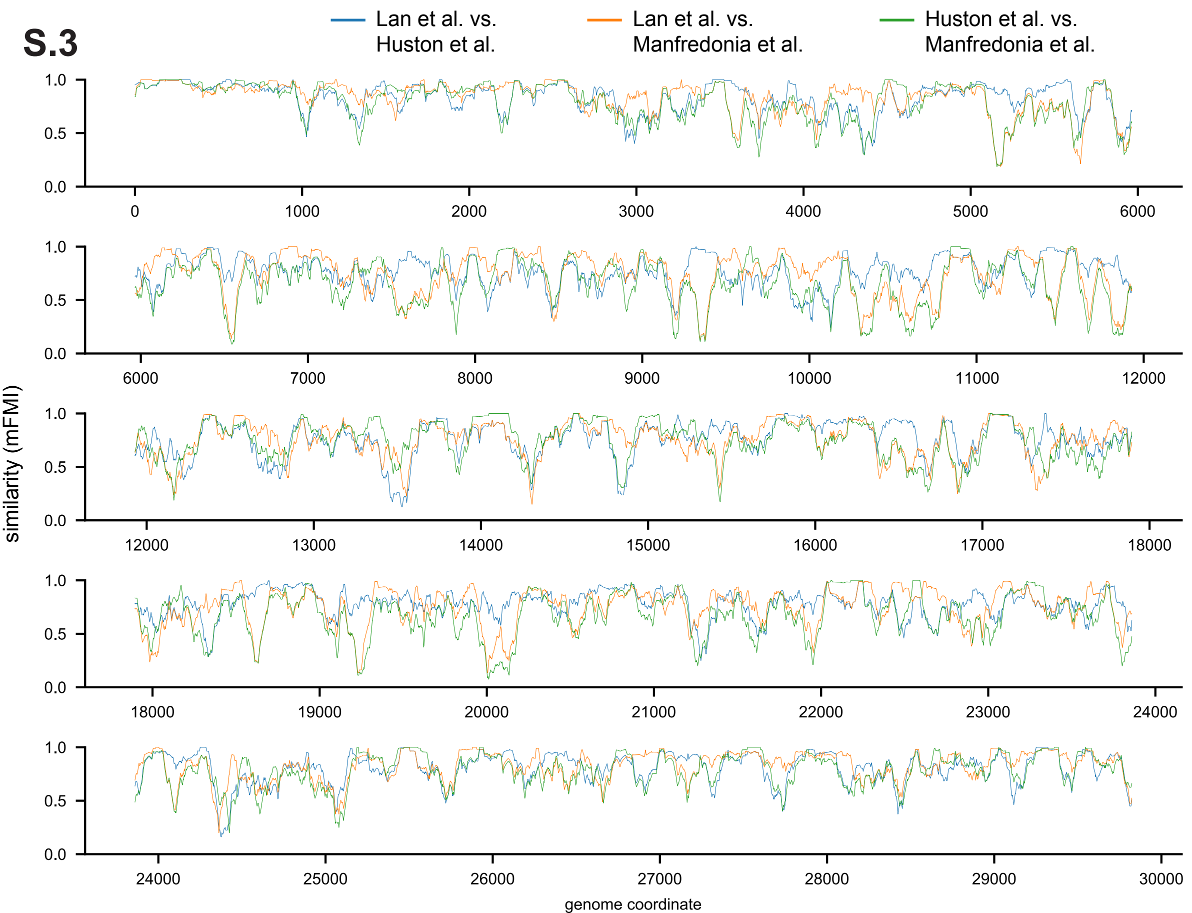
Lan et al. vs. Huston et al.
Lan et al. vs. Manfredonia et al.
Huston et al. vs. Manfredonia et al.
similarity (mFMI)
genome coordinate
1.0
0.5
0.0
0
1000
2000
3000
4000
5000
6000
7000
8000
9000
10000
11000
12000
13000
14000
15000
16000
17000
18000
19000
20000
21000
22000
23000
24000
25000
26000
27000
28000
29000
30000

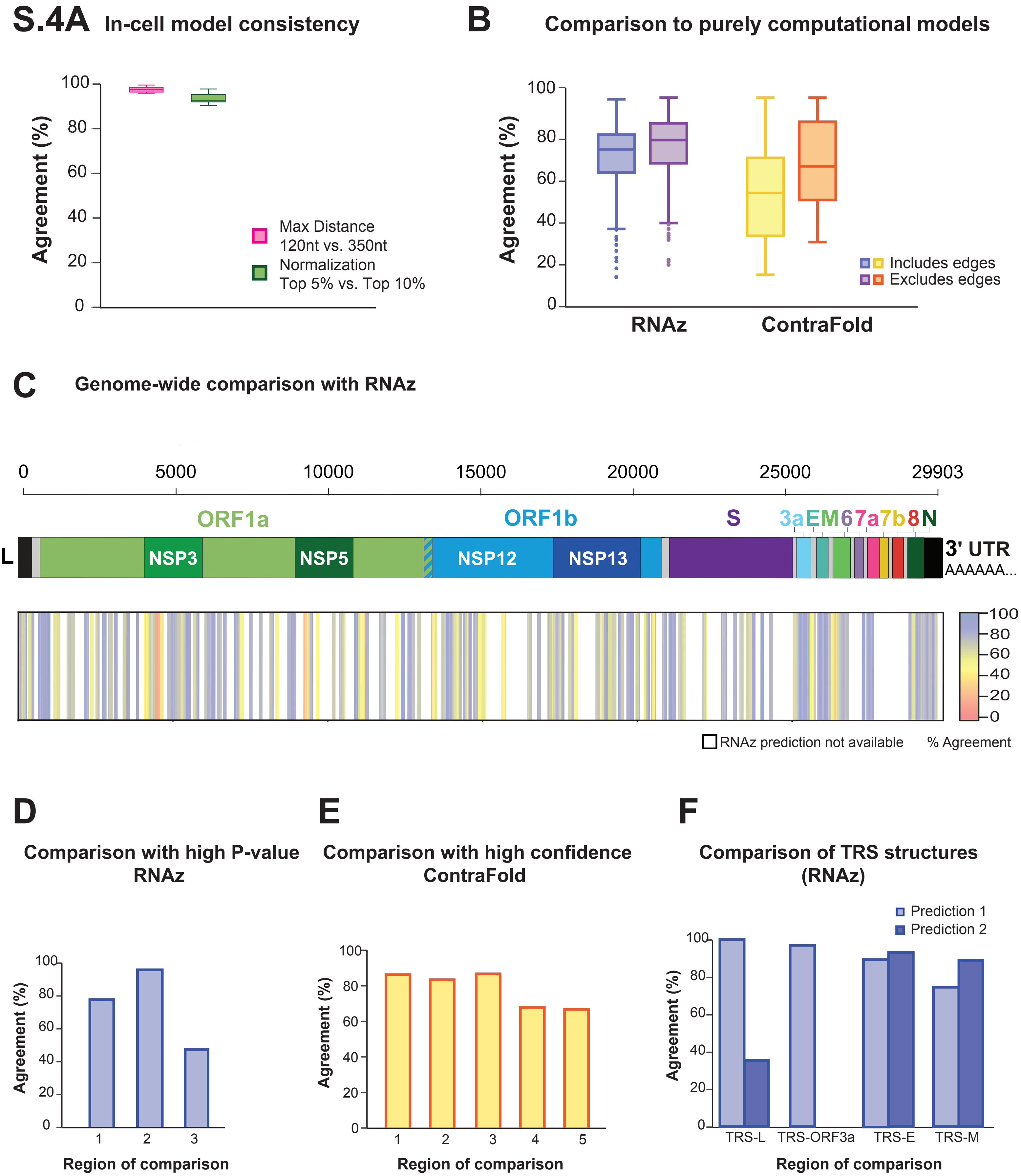
S.4A In-cell model consistency
Agreement (%)
100
80
60
40
20
0
Max Distance
120nt vs. 350nt
Normalization
Top 5% vs. Top 10%
B Comparison to purely computational models
Agreement (%)
100
80
60
40
20
0
Includes edges
Excludes edges
RNAz
ContraFold
C Genome-wide comparison with RNAz
0
5000
10000
15000
20000
25000
29903
ORF1a
ORF1b
S
3a E M 6 7a 7b 8 N
L
NSP3
NSP5
NSP12
NSP13
3' UTR
AAAAAA...
100
80
60
40
20
0
RNAz prediction not available
% Agreement
D
Comparison with high P-value RNAz
Agreement (%)
100
80
60
40
20
0
1
2
3
Region of comparison
E
Comparison with high confidence ContraFold
Agreement (%)
100
80
60
40
20
0
1
2
3
4
5
Region of comparison
F
Comparison of TRS structures (RNAz)
Prediction 1
Prediction 2
Agreement (%)
100
80
60
40
20
0
TRS-L
TRS-ORF3a
TRS-E
TRS-M
Region of comparison

# S.5A

### Structured regions and accessible regions across the genome

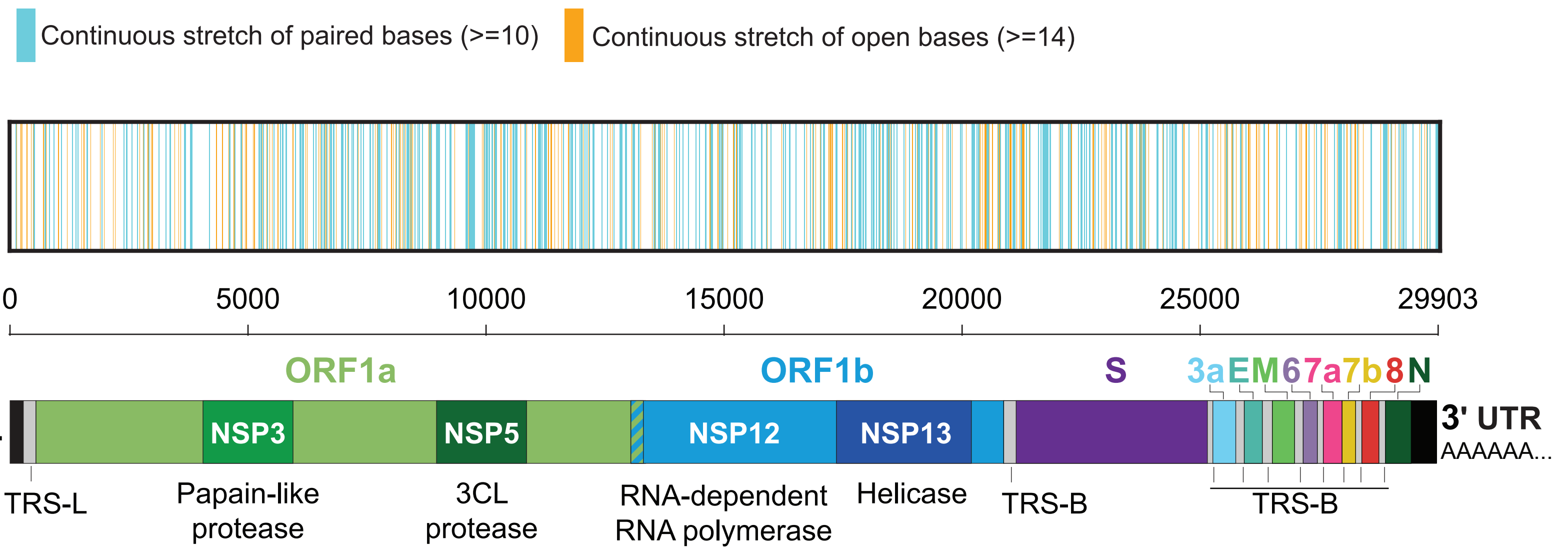


### B Structures at the TRS-L and TRS-B

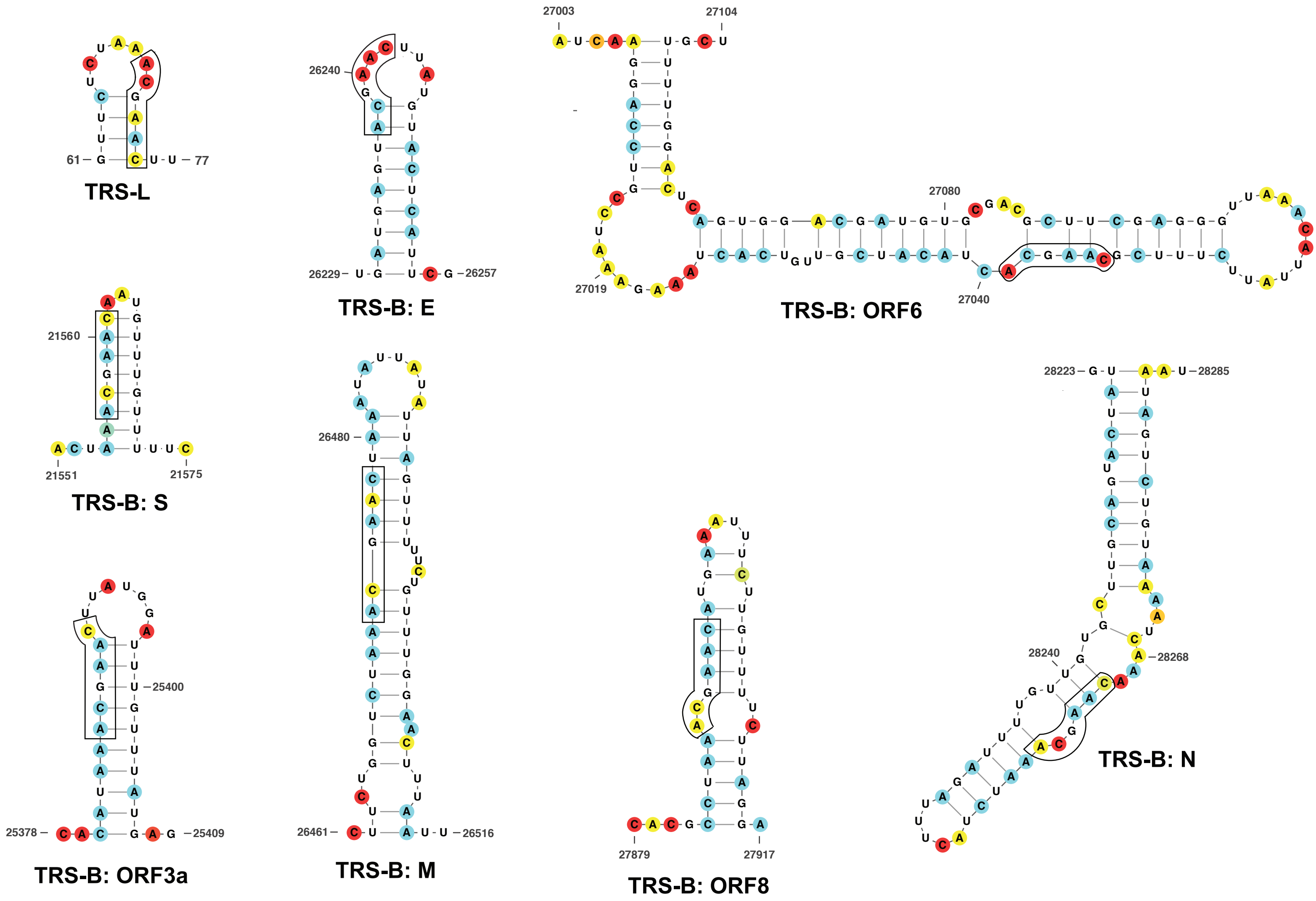

S.6

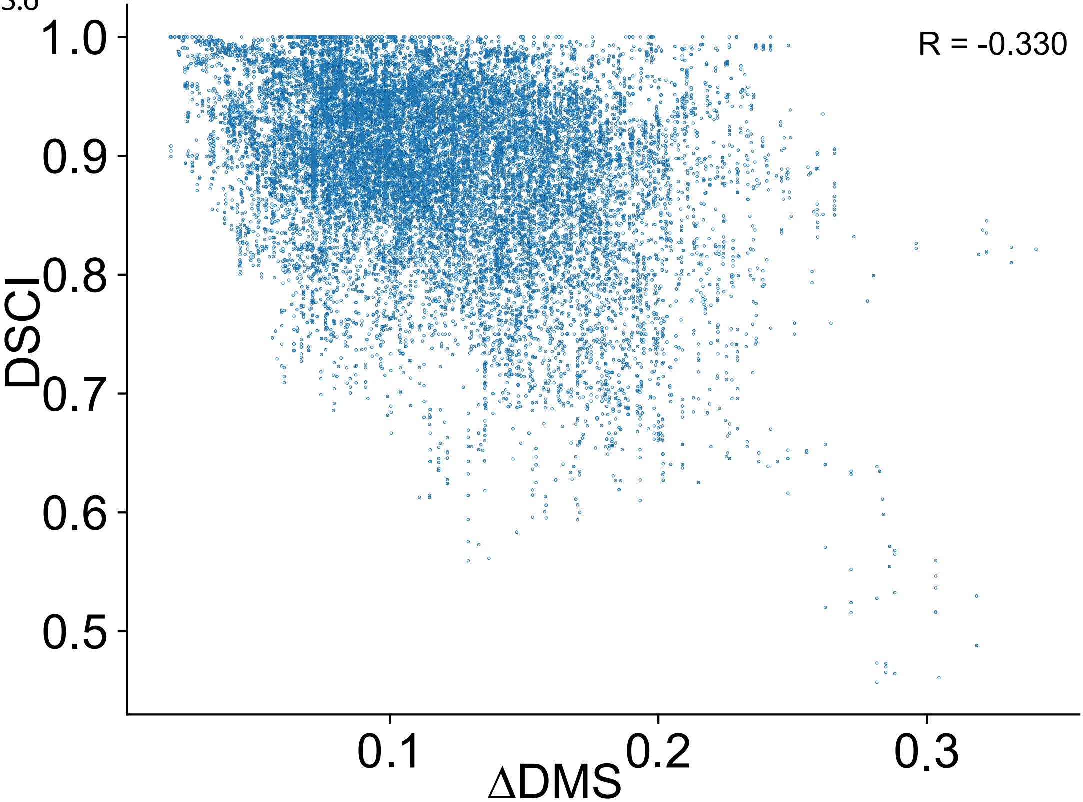

**S. Table 1`**

| index | start | end | length |
|---|---|---|---|
| 1 | 128 | 149 | 22 |
| 2 | 505 | 528 | 24 |
| 3 | 726 | 756 | 31 |
| 4 | 823 | 837 | 15 |
| 5 | 926 | 939 | 14 |
| 6 | 1071 | 1089 | 19 |
| 7 | 1304 | 1321 | 18 |
| 8 | 1330 | 1359 | 30 |
| 9 | 1377 | 1399 | 23 |
| 10 | 1421 | 1438 | 18 |
| 11 | 1573 | 1598 | 26 |
| 12 | 1610 | 1636 | 27 |
| 13 | 1919 | 1939 | 21 |
| 14 | 2385 | 2398 | 14 |
| 15 | 2435 | 2458 | 24 |
| 16 | 2550 | 2568 | 19 |
| 17 | 2667 | 2682 | 16 |
| 18 | 2801 | 2819 | 19 |
| 19 | 2901 | 2918 | 18 |
| 20 | 3116 | 3129 | 14 |
| 21 | 3258 | 3273 | 16 |
| 22 | 3349 | 3371 | 23 |
| 23 | 3647 | 3662 | 16 |
| 24 | 3668 | 3683 | 16 |
| 25 | 3733 | 3818 | 86 |
| 26 | 3863 | 3883 | 21 |
| 27 | 4101 | 4122 | 22 |
| 28 | 4264 | 4307 | 44 |
| 29 | 4580 | 4608 | 29 |
| 30 | 4619 | 4645 | 27 |
| 31 | 4702 | 4715 | 14 |
| 32 | 4826 | 4846 | 21 |
| 33 | 4877 | 4897 | 21 |
| 34 | 5020 | 5036 | 17 |
| 35 | 5297 | 5311 | 15 |
| 36 | 5382 | 5399 | 18 |
| 37 | 5522 | 5535 | 14 |
| 38 | 5647 | 5662 | 16 |
| 39 | 5688 | 5715 | 28 |
| 40 | 5773 | 5795 | 23 |
| 41 | 5933 | 5949 | 17 |
| 42 | 5955 | 5969 | 15 |

| | | | |
|---|---|---|---|
| 43 | 6058 | 6090 | 33 |
| 44 | 6103 | 6145 | 43 |
| 45 | 6307 | 6325 | 19 |
| 46 | 6346 | 6362 | 17 |
| 47 | 6390 | 6403 | 14 |
| 48 | 6505 | 6532 | 28 |
| 49 | 6538 | 6554 | 17 |
| 50 | 6567 | 6589 | 23 |
| 51 | 6622 | 6640 | 19 |
| 52 | 6697 | 6736 | 40 |
| 53 | 6771 | 6785 | 15 |
| 54 | 6925 | 6973 | 49 |
| 55 | 7010 | 7025 | 16 |
| 56 | 7073 | 7092 | 20 |
| 57 | 7146 | 7193 | 48 |
| 58 | 7214 | 7229 | 16 |
| 59 | 7260 | 7280 | 21 |
| 60 | 7391 | 7404 | 14 |
| 61 | 7511 | 7536 | 26 |
| 62 | 7547 | 7567 | 21 |
| 63 | 7590 | 7603 | 14 |
| 64 | 7690 | 7716 | 27 |
| 65 | 7725 | 7743 | 19 |
| 66 | 8075 | 8095 | 21 |
| 67 | 8128 | 8143 | 16 |
| 68 | 8290 | 8327 | 38 |
| 69 | 8336 | 8349 | 14 |
| 70 | 8379 | 8392 | 14 |
| 71 | 8459 | 8472 | 14 |
| 72 | 8523 | 8537 | 15 |
| 73 | 8556 | 8576 | 21 |
| 74 | 8621 | 8647 | 27 |
| 75 | 8669 | 8690 | 22 |
| 76 | 8766 | 8779 | 14 |
| 77 | 8800 | 8822 | 23 |
| 78 | 8906 | 8942 | 37 |
| 79 | 8957 | 8992 | 36 |
| 80 | 9096 | 9117 | 22 |
| 81 | 9131 | 9158 | 28 |
| 82 | 9187 | 9224 | 38 |
| 83 | 9281 | 9294 | 14 |
| 84 | 9527 | 9546 | 20 |
| 85 | 9555 | 9569 | 15 |

| | | | |
|---|---|---|---|
| 86 | 9624 | 9680 | 57 |
| 87 | 9913 | 9931 | 19 |
| 88 | 10006 | 10030 | 25 |
| 89 | 10067 | 10096 | 30 |
| 90 | 10157 | 10174 | 18 |
| 91 | 10440 | 10456 | 17 |
| 92 | 10500 | 10536 | 37 |
| 93 | 10551 | 10614 | 64 |
| 94 | 10754 | 10781 | 28 |
| 95 | 10912 | 10927 | 16 |
| 96 | 11040 | 11082 | 43 |
| 97 | 11118 | 11131 | 14 |
| 98 | 11165 | 11230 | 66 |
| 99 | 11560 | 11577 | 18 |
| 100 | 11728 | 11742 | 15 |
| 101 | 11755 | 11772 | 18 |
| 102 | 11814 | 11830 | 17 |
| 103 | 11910 | 11928 | 19 |
| 104 | 11957 | 11970 | 14 |
| 105 | 12014 | 12027 | 14 |
| 106 | 12055 | 12129 | 75 |
| 107 | 12173 | 12220 | 48 |
| 108 | 12281 | 12309 | 29 |
| 109 | 12314 | 12330 | 17 |
| 110 | 12484 | 12498 | 15 |
| 111 | 12619 | 12635 | 17 |
| 112 | 12665 | 12696 | 32 |
| 113 | 12741 | 12766 | 26 |
| 114 | 12775 | 12791 | 17 |
| 115 | 12881 | 12928 | 48 |
| 116 | 12967 | 12988 | 22 |
| 117 | 13052 | 13065 | 14 |
| 118 | 13327 | 13347 | 21 |
| 119 | 13723 | 13736 | 14 |
| 120 | 13912 | 13957 | 46 |
| 121 | 13990 | 14005 | 16 |
| 122 | 14056 | 14070 | 15 |
| 123 | 14170 | 14185 | 16 |
| 124 | 14405 | 14418 | 14 |
| 125 | 14642 | 14690 | 49 |
| 126 | 14819 | 14841 | 23 |
| 127 | 14861 | 14889 | 29 |
| 128 | 14900 | 14915 | 16 |

| | | | |
|---|---|---|---|
| 129 | 14926 | 14942 | 17 |
| 130 | 15139 | 15156 | 18 |
| 131 | 15196 | 15213 | 18 |
| 132 | 15236 | 15272 | 37 |
| 133 | 15459 | 15472 | 14 |
| 134 | 15498 | 15514 | 17 |
| 135 | 15560 | 15573 | 14 |
| 136 | 15695 | 15720 | 26 |
| 137 | 15801 | 15818 | 18 |
| 138 | 15902 | 15918 | 17 |
| 139 | 16199 | 16213 | 15 |
| 140 | 16297 | 16312 | 16 |
| 141 | 16419 | 16434 | 16 |
| 142 | 16592 | 16607 | 16 |
| 143 | 16712 | 16737 | 26 |
| 144 | 16761 | 16775 | 15 |
| 145 | 16787 | 16805 | 19 |
| 146 | 16938 | 16954 | 17 |
| 147 | 16990 | 17026 | 37 |
| 148 | 17273 | 17294 | 22 |
| 149 | 17491 | 17504 | 14 |
| 150 | 17515 | 17538 | 24 |
| 151 | 17561 | 17593 | 33 |
| 152 | 17651 | 17667 | 17 |
| 153 | 17739 | 17796 | 58 |
| 154 | 17822 | 17853 | 32 |
| 155 | 17875 | 17898 | 24 |
| 156 | 18008 | 18052 | 45 |
| 157 | 18102 | 18115 | 14 |
| 158 | 18151 | 18180 | 30 |
| 159 | 18219 | 18260 | 42 |
| 160 | 18321 | 18360 | 40 |
| 161 | 18371 | 18384 | 14 |
| 162 | 18415 | 18436 | 22 |
| 163 | 18471 | 18486 | 16 |
| 164 | 18531 | 18546 | 16 |
| 165 | 18789 | 18802 | 14 |
| 166 | 18918 | 18937 | 20 |
| 167 | 18965 | 18991 | 27 |
| 168 | 19003 | 19018 | 16 |
| 169 | 19046 | 19065 | 20 |
| 170 | 19249 | 19271 | 23 |
| 171 | 19569 | 19593 | 25 |

| | | | |
|---|---|---|---|
| 172 | 19601 | 19615 | 15 |
| 173 | 19623 | 19664 | 42 |
| 174 | 19861 | 19876 | 16 |
| 175 | 19883 | 19899 | 17 |
| 176 | 20003 | 20023 | 21 |
| 177 | 20092 | 20161 | 70 |
| 178 | 20198 | 20213 | 16 |
| 179 | 20230 | 20253 | 24 |
| 180 | 20403 | 20443 | 41 |
| 181 | 20652 | 20667 | 16 |
| 182 | 20693 | 20708 | 16 |
| 183 | 20794 | 20835 | 42 |
| 184 | 20947 | 21008 | 62 |
| 185 | 21123 | 21147 | 25 |
| 186 | 21235 | 21248 | 14 |
| 187 | 21304 | 21317 | 14 |
| 188 | 21478 | 21501 | 24 |
| 189 | 21573 | 21752 | 180 |
| 190 | 21855 | 21868 | 14 |
| 191 | 21949 | 21974 | 26 |
| 192 | 22115 | 22137 | 23 |
| 193 | 22153 | 22170 | 18 |
| 194 | 22390 | 22405 | 16 |
| 195 | 22442 | 22459 | 18 |
| 196 | 22547 | 22573 | 27 |
| 197 | 22611 | 22624 | 14 |
| 198 | 22736 | 22750 | 15 |
| 199 | 23010 | 23047 | 38 |
| 200 | 23079 | 23135 | 57 |
| 201 | 23155 | 23168 | 14 |
| 202 | 23306 | 23324 | 19 |
| 203 | 23395 | 23409 | 15 |
| 204 | 23414 | 23460 | 47 |
| 205 | 23590 | 23607 | 18 |
| 206 | 23626 | 23641 | 16 |
| 207 | 23655 | 23695 | 41 |
| 208 | 23705 | 23725 | 21 |
| 209 | 23910 | 23934 | 25 |
| 210 | 23960 | 23973 | 14 |
| 211 | 23984 | 23999 | 16 |
| 212 | 24095 | 24108 | 14 |
| 213 | 24165 | 24184 | 20 |
| 214 | 24235 | 24250 | 16 |

| | | | |
|---|---|---|---|
| 215 | 24270 | 24304 | 35 |
| 216 | 24337 | 24353 | 17 |
| 217 | 24361 | 24384 | 24 |
| 218 | 24490 | 24503 | 14 |
| 219 | 24633 | 24647 | 15 |
| 220 | 24888 | 24911 | 24 |
| 221 | 24985 | 25008 | 24 |
| 222 | 25015 | 25032 | 18 |
| 223 | 25133 | 25148 | 16 |
| 224 | 25408 | 25439 | 32 |
| 225 | 25546 | 25563 | 18 |
| 226 | 25579 | 25592 | 14 |
| 227 | 25703 | 25719 | 17 |
| 228 | 25798 | 25816 | 19 |
| 229 | 25823 | 25836 | 14 |
| 230 | 26198 | 26224 | 27 |
| 231 | 26508 | 26531 | 24 |
| 232 | 26722 | 26742 | 21 |
| 233 | 26862 | 26886 | 25 |
| 234 | 26902 | 26919 | 18 |
| 235 | 27128 | 27148 | 21 |
| 236 | 27289 | 27302 | 14 |
| 237 | 27363 | 27408 | 46 |
| 238 | 27546 | 27560 | 15 |
| 239 | 27669 | 27692 | 24 |
| 240 | 27751 | 27777 | 27 |
| 241 | 27800 | 27828 | 29 |
| 242 | 27869 | 27882 | 14 |
| 243 | 27997 | 28012 | 16 |
| 244 | 28142 | 28163 | 22 |
| 245 | 28333 | 28346 | 14 |
| 246 | 28431 | 28455 | 25 |
| 247 | 28688 | 28711 | 24 |
| 248 | 28721 | 28754 | 34 |
| 249 | 28767 | 28781 | 15 |
| 250 | 29004 | 29025 | 22 |
| 251 | 29074 | 29089 | 16 |
| 252 | 29110 | 29133 | 24 |
| 253 | 29155 | 29178 | 24 |
| 254 | 29307 | 29320 | 14 |
| 255 | 29488 | 29502 | 15 |
| 256 | 29614 | 29628 | 15 |
| 257 | 29658 | 29673 | 16 |

| | | | |
|---|---|---|---|
| 258 | 29786 | 29800 | 15 |
| 259 | 29824 | 29844 | 21 |
